# Non-homologous end joining shapes the genomic rearrangement landscape of chromothripsis from mitotic errors

**DOI:** 10.1101/2023.08.10.552800

**Authors:** Qing Hu, Jose Espejo Valle-Inclan, Rashmi Dahiya, Alison Guyer, Alice Mazzagatti, Elizabeth G. Maurais, Justin L. Engel, Isidro Cortés-Ciriano, Peter Ly

## Abstract

Errors in mitosis can generate micronuclei that entrap mis-segregated chromosomes, which are susceptible to catastrophic fragmentation through a process termed chromothripsis. The reassembly of fragmented chromosomes by error-prone DNA double-strand break (DSB) repair generates a spectrum of simple and complex genomic rearrangements that are associated with human cancers and disorders. How specific DSB repair pathways recognize and process these lesions remains poorly understood. Here we used CRISPR/Cas9 to systematically inactivate distinct DSB processing or repair pathways and interrogated the rearrangement landscape of fragmented chromosomes from micronuclei. Deletion of canonical non-homologous end joining (NHEJ) components, including DNA-PKcs, LIG4, and XLF, substantially reduced the formation of complex rearrangements and shifted the rearrangement landscape toward simple alterations without the characteristic patterns of cancer-associated chromothripsis. Following reincorporation into the nucleus, fragmented chromosomes localize within micronuclei bodies (MN bodies) and undergo successful ligation by NHEJ within a single cell cycle. In the absence of NHEJ, chromosome fragments were rarely engaged by polymerase theta-mediated alternative end-joining or recombination-based mechanisms, resulting in delayed repair kinetics and persistent 53BP1-labeled MN bodies in the interphase nucleus. Prolonged DNA damage signaling from unrepaired fragments ultimately triggered cell cycle arrest. Thus, we provide evidence supporting NHEJ as the exclusive DSB repair pathway generating complex rearrangements following chromothripsis from mitotic errors.

## INTRODUCTION

Chromosome segregation during mitosis must be accurately executed to maintain genome stability. Mitotic errors can result in the formation of aberrant nuclear structures called micronuclei that entrap mis-segregated chromosomes or chromosome arms. The irreversible rupture of the micronuclear envelope^1,2^ triggers the loss of nucleocytoplasmic compartmentalization and the acquisition of DNA double-strand breaks (DSBs)^1,3–6^. Extensive DSBs can promote the catastrophic fragmentation of the micronucleated chromosome during the subsequent mitosis^4,7^, a process that has been termed chromothripsis^8^. Fragmented chromosomes remain spatially clustered during mitosis and reincorporate into the nucleus of one or both daughter cell(s), manifesting during interphase as sub-nuclear territories known as ‘micronuclei bodies’ (MN bodies) that engage the DNA damage response^4,9–11^.

Chromothripsis frequently drives complex and localized genomic rearrangements owing to the error-prone reassembly of the fragmented chromosome^12^. These rearrangements are common across diverse cancer types^8,13,14^ and can be characterized by seemingly random structural variants that are clustered along one or a few chromosome(s)^15^. In addition to complex rearrangements, a diverse spectrum of chromosomal abnormalities can be generated from micronuclei formation, including simple arm-level deletions, insertions, and translocations^5,16^. Based on the sequence features at rearrangement breakpoint junctions, several DSB repair pathways have been predicted to underlie the formation of complex rearrangements following chromothripsis^5,13,16^.

Multiple DSB repair pathways are operative in mammalian cells to process detrimental DNA lesions. The canonical non-homologous end joining (NHEJ) pathway is active throughout the cell cycle and directly ligates two DSB ends through the recruitment of Ku70/80 and activity of DNA-PKcs, XLF, and DNA ligase 4–XRCC4^17^. Alternative end-joining (alt-EJ) occurs independently of canonical NHEJ factors, with DNA polymerase theta (Polθ)-dependent microhomology-mediated end joining accounting for most known alt-EJ events^18^. Whereas NHEJ ligates DSBs without homology, alt-EJ relies on the resection of DSB ends to expose short stretches of homologous sequence – known as microhomology – at the repair junction. Both homologous recombination (HR) and single-strand annealing (SSA) require more extensive DNA end resection to generate extended 3′ single-strand DNA (ssDNA) tails. HR is most active following genome duplication in S-phase, a period when ssDNA tails can invade a homologous DNA sequence (e.g., a sister chromatid) and uses it as a template for synthesis to repair the DSB, which involves BRCA1, BRCA2, RAD51, and RAD54^19^. SSA requires annealing of homologous repeats to form the synapsis intermediate before ligation, a process that is mediated by RAD52^20^. The DSB repair pathways described can be mutagenic when multiple DSBs are present and/or incorrect sequences are used for recombination^21–25^.

In cancers genomes and germline disorders with chromothripsis, the majority of rearrangement breakpoints harbor blunt-ended junctions without homology; however, microhomology signatures – which can be arbitrarily defined to include as little as one nucleotide of homology – have also been reported^8,13,14,26–30^. Although similar observations have been described in experimental models of chromothripsis^5,16,31^, the extent to which specific DSB repair pathways contribute to reassembling fragmented chromosomes from micronuclei has not been systematically characterized. We previously showed that depletion of DNA PKcs-and DNA ligase 4, two important components of NHEJ, was sufficient to reduce, but not completely abrogate, the formation of rearrangements from micronuclei^7,16^. This approach was limited by a partial reduction of key genes by RNA interference, including DNA repair enzymes whose activity may remain functional at low levels. Additionally, chromothripsis from DSB repair-deficient murine tumors^32^ and human cells escaping telomere crisis^33^ appear to arise independently of NHEJ. It thus remains unclear how DNA lesions from micronuclei are processed and which DSB repair pathway(s) can generate the range of simple and complex rearrangements that are pervasive in cancer genomes.

To determine how specific DSB repair pathways shape the rearrangement landscape of mitotic errors, we leveraged a strategy termed CEN-SELECT, which enables the controlled induction of micronuclei containing the Y chromosome harboring a neomycin-resistance (*neo^R^*) marker^7,16^. Using this approach, exposure to doxycycline and auxin (DOX/IAA) induces the replacement of the centromeric histone H3 variant CENP-A with a chimeric mutant that functionally inactivates the Y centromere. Following mitotic mis-segregation into micronuclei and chromosome fragmentation, selection for the Y-encoded *neo^R^* marker allows for the isolation of a diverse spectrum of rearrangement types^16^. By generating a series of gene deletions spanning each DSB repair pathway, here we identified the NHEJ pathway as the predominant repair mechanism in forming complex rearrangements following chromothripsis. In the absence of NHEJ, chromosome fragments were rarely reassembled by non-NHEJ DSB repair pathways, resulting in persistent DNA damage within the nucleus as MN bodies that triggered cell cycle arrest.

## RESULTS

### NHEJ is the primary DSB repair pathway for fragmented chromosomes from micronuclei

To study the contributions of each DSB repair pathway to chromothripsis, we first generated isogenic DLD-1 knockout (KO) cells in the background of the CEN-SELECT system (**Figure 1a**). This was achieved by delivering Cas9 ribonucleoproteins (RNPs) in complex with sgRNAs targeting eight genes spanning multiple DSB repair-related processes, including canonical NHEJ (*PRKDC*, *LIG4*, *NHEJ1*), DNA end protection (*TP53BP1*), alt-EJ (*POLQ*), SSA (*RAD52*), HR (*RAD54L*), and DNA end resection (*NBN*). In addition to their critical function in a specific DSB repair pathway, these non-essential genes were selected because cells are able to maintain a mostly diploid karyotype in its absence. Gene KOs were created using either a single sgRNA to induce insertion/deletion mutations or dual sgRNAs to generate frameshift deletions that are discernable by polymerase chain reaction (PCR) from the bulk cell population and single cell-derived clones (**Extended Data Figure 1a**). Twenty-two clones harboring biallelic inactivation of the target genes were confirmed by PCR, Sanger sequencing, and/or immunoblotting (**Extended Data Figure 1b-c**). Following DOX/IAA treatment, all KO clones generated micronuclei at a frequency comparable to WT cells (**Extended Data Figure 1d-e**), which resulted in the shattering of the Y chromosome that can be detected by DNA fluorescence *in situ* hybridization (FISH) on metaphase spreads (**Extended Data Figure 1f-g**).

**Figure 1.**
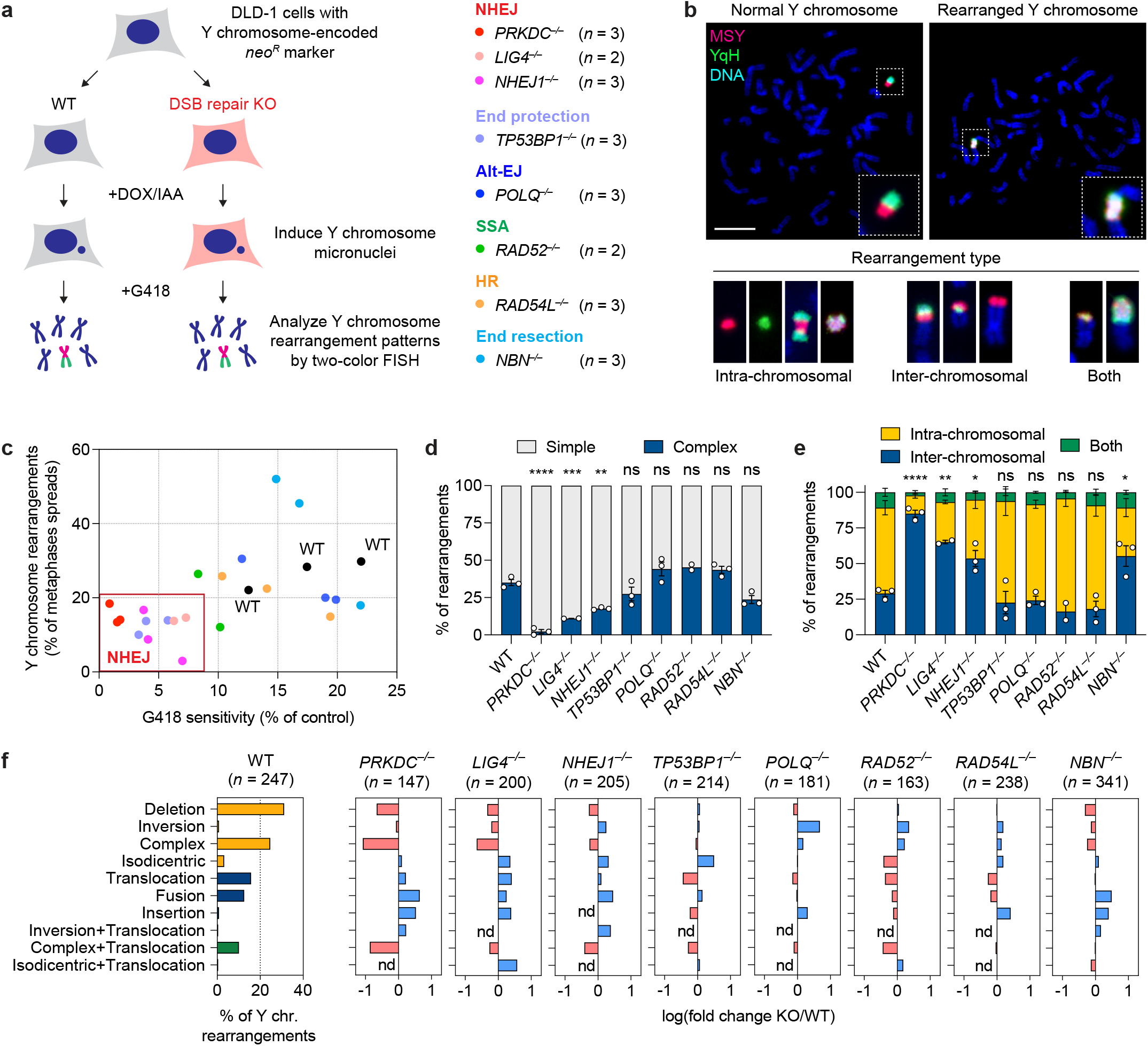
Genomic rearrangement landscape of mis-segregated chromosomes in the absence of specific DNA double-strand break (DSB) repair pathways. **a)** Experimental approach to survey the impact of specific DSB repair pathways on chromosome rearrangements induced by micronucleus formation. Biallelic gene knockouts (KOs) were generated in the background of the CEN-SELECT system in isogenic DLD-1 cells. Y chromosome-specific mis-segregation into micronuclei and rearrangements were induced by treatment with doxycycline and auxin (DOX/IAA). b) Representative examples of metaphase spreads with normal or derivative Y chromosomes. Different types of rearrangements can be visualized by DNA fluorescence *in situ* hybridization (FISH) using probes targeting the euchromatic portion of the male-specific region (MSY, red) and the heterochromatic region (YqH, green) of the Y chromosome. Rearrangements were induced by 3d DOX/IAA treatment followed by G418 selection. Scale bar, 10 µm. c) Plot summarizing the effect on cell viability after G418 selection (*x-axis*) and rearrangement frequency of the Y chromosome (*y-axis*) for each DSB repair KO clone. d) Proportion of Y chromosomes exhibiting simple or complex rearrangements, as determined by metaphase FISH, following transient centromere inactivation. e) Proportion of inter-and/or intra-chromosomal rearrangements. Data in (d) and (e) represent the mean ± SEM of *n* = 3 independent experiments for WT, *n* = 2 KO clones for *LIG4* and *RAD52*, and *n* = 3 KO clones for *PRKDC*, *NHEJ1*, *TP53BP1*, *POLQ*, *RAD54L*, and *NBN*; statistical analyses were calculated by ordinary one-way ANOVA test with multiple comparisons. ns, not significant; \**P* ≤ 0.05; \*\**P* ≤ 0.01; \*\*\**P* ≤ 0.001; \*\*\*\**P* ≤ 0.0001. f) Left: Distribution of Y chromosome rearrangement types as determined by metaphase FISH following 3d DOX/IAA treatment and G418 selection. Data are pooled from 3 independent experiments. Right: Plots depict the mean fold change in each rearrangement type as compared to WT cells. Sample sizes indicate the number of rearranged Y chromosomes examined; data are pooled from 2 or 3 individual KO clones per gene. See also Extended Data Figure 2.

Following Y chromosome shattering, the reassembly of the *neo^R^*-containing fragment into a stable derivative chromosome confers long-term resistance to G418 selection and produces rearrangements that can be visualized by cytogenetics (**Figure 1b**). Cells that cannot maintain the *neo^R^* fragment are rendered sensitive to G418 selection^16^. Among the 22 KO clones generated, NHEJ-deficient cells lacking either DNA-PKcs, DNA ligase 4, or XLF exhibited decreased survival in G418, indicative of their failure to maintain a functional *neo^R^* marker after micronucleation and fragmentation of the Y chromosome (**Figure 1c**). Loss of 53BP1, which promotes NHEJ by protecting DSB ends from undergoing resection, similarly resulted in decreased G418 survival (**Figure 1c**). We then compared rearrangement frequencies across the surviving fraction of cells by using two DNA paint probes targeting each half of the Y chromosome to visualize a range of Y chromosome-specific rearrangements (**Figure 1b**). NHEJ-deficient cells surviving G418 selection showed an overall decrease in rearrangement frequencies regardless of rearrangement type (**Figure 1c**). In contrast, deletion of *POLQ*, *RAD52*, *RAD54L*, and *NBN* – which are involved in alt-EJ, SSA, HR, and DNA end-resection, respectively – had minimal to no effect on both cell survival under G418 selection and rearrangement frequencies following the induction of Y chromosome micronucleation (**Figure 1c**).

### Genomic rearrangement landscape of DSB repair deficiency

We next examined how loss of a specific DSB repair pathway influences the spectrum of rearrangement types generated from micronuclei formation. In WT cells, the induction of Y chromosome mis-segregation followed by G418 selection for retention of *neo^R^* generated a diverse range of simple and complex intra-and inter-chromosomal rearrangements (**Figure 1d-f**, **Extended Data Figure 2**), consistent with prior studies^16^. In the absence of NHEJ, however, the rearrangement landscape shifted towards relatively simple inter-chromosomal rearrangements, which were largely comprised of non-reciprocal translocations and whole-chromosome fusions (**Figure 1d-f**, **Extended Data Figure 2**). Notably, there was a sharp reduction in complex rearrangements (**Figure 1d**), which were distinguishable by the co-localization of the two-colored FISH probes that are normally separated on metaphase Y chromosomes (**Figure 1b**). We previously showed that this cytogenetics-based approach is highly concordant with the use of whole-genome sequencing (WGS) to call chromothripsis events^16^. Reduced intra-chromosomal and complex rearrangements were unexpectedly observed in cells lacking Nbs1, a component of the MRN complex that resects DSB ends^34^, but not in *TP53BP1*, *POLQ*, *RAD52* and *RAD54L* KO cells (**Figure 1d-f**, **Extended Data Figure 2**).

To determine which DSB repair pathways are required to produce stable and genetically heritable derivative chromosomes from micronuclei, the KO panel of cells were cultured under sustained centromere inactivation and continuous G418 selection over the span of approximately 30 days (**Extended Data Figure 3a**). In agreement with transient centromere inactivation, WT cells and cells deficient in either the alt-EJ, HR, or SSA pathways exhibited reduced growth rates during early passages followed by a recovery over time (**Extended Data Figure 3b**), reflecting cell death owing to the loss of the *neo^R^* marker upon the induction of Y chromosome mis-segregation and the subsequent formation of stable Y chromosome rearrangements^16^. By contrast, NHEJ-deficient cells exhibited a further reduction in growth rate during early passages that was accompanied by delayed recovery in proliferation (**Extended Data Figure 3b**). A similar trend was observed in cells lacking *NBN*, suggesting a possible contribution by factors that promote the resection of DSB ends. In agreement with reduced growth in G418, metaphase FISH revealed that NHEJ-deficient cells harbored fewer derivative Y chromosomes, which consisted mostly of simple rearrangements (**Figure 2a-b**, **Extended Data Figure 3c**).

**Figure 2.**
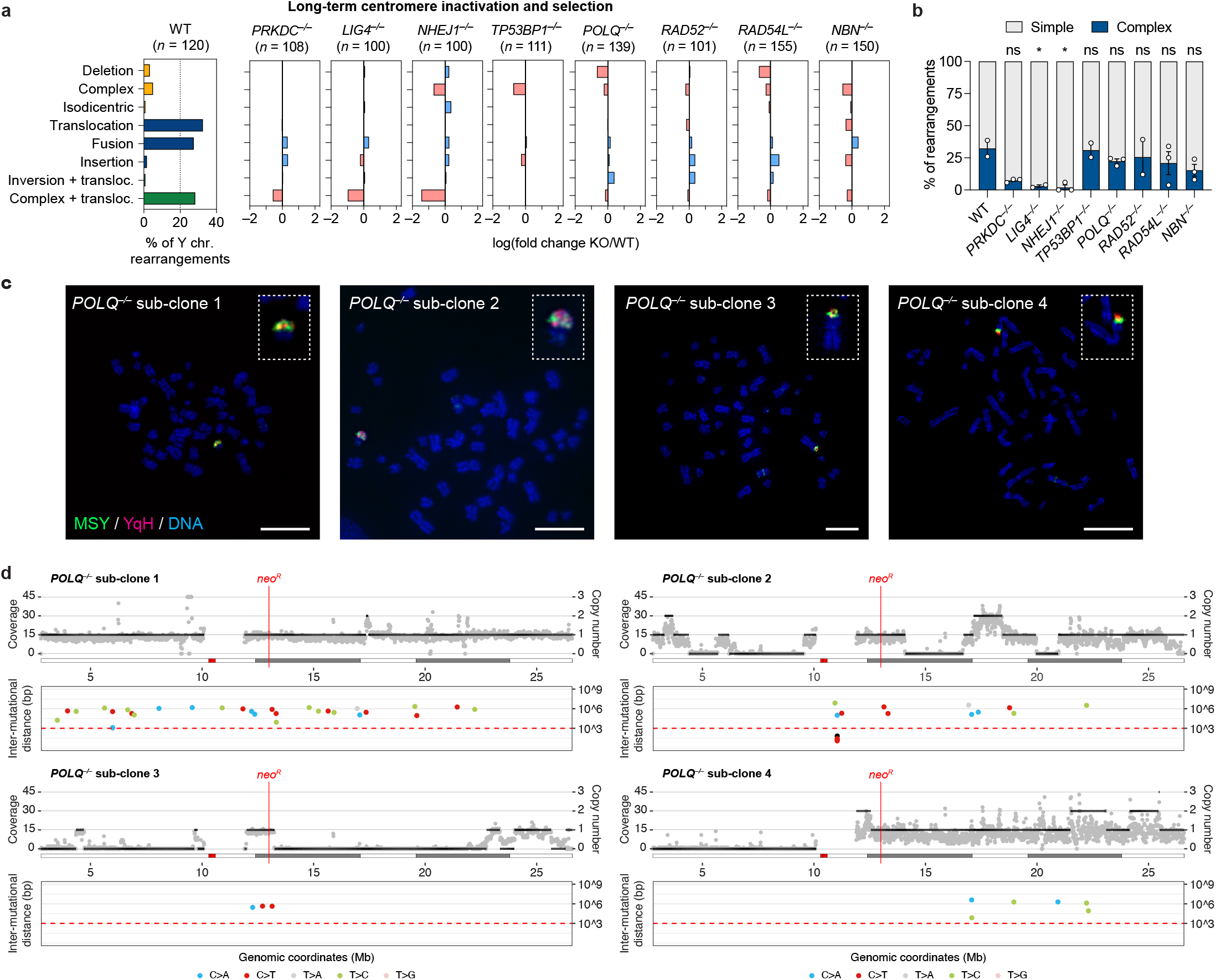
DSB repair pathways beyond NHEJ are dispensable for generating complex rearrangements from micronuclei. **a)** Left: Distribution of Y chromosome rearrangement types as determined by metaphase FISH following continuous passage in DOX/IAA and G418 for ∼30 days. Data are pooled from 2 independent experiments. Right: Plots depict the mean fold change in each rearrangement type as compared to WT cells. Sample sizes indicate the number of rearranged Y chromosomes examined; data are pooled from 2 or 3 individual KO clones per gene. See also **Extended Data Figure 3c**. **b)** Proportion of Y chromosomes exhibiting simple or complex rearrangements, as determined by metaphase FISH, following sustained centromere inactivation. Data represent the mean ± SEM of *n* = 2 independent experiments for WT, *n* = 2 KO clones for *LIG4* and *RAD52*, and *n* = 3 KO clones for *PRKDC*, *NHEJ1*, *TP53BP1*, *POLQ*, *RAD54L*, and *NBN*; statistical analyses were calculated by ordinary one-way ANOVA test with multiple comparisons. ns, not significant; \**P* ≤ 0.05. **c)** Cytogenetic characterization of *POLQ* KO sub-clones harboring complex Y chromosome rearrangements following sustained centromere inactivation. Scale bar, 10 µm. **d)** Whole-genome sequencing analyses of *POLQ* KO sub-clones with complex Y chromosome rearrangements exhibiting oscillating DNA copy-number patterns. For each subclone, sequencing depth (grey dots), copy number information (black lines; top), and inter-mutational distances (bottom) for the mappable regions of the Y chromosome are shown.

### NHEJ-deficient cells fail to generate complex rearrangements from micronuclei

A proportion of chromothriptic breakpoint junctions in tumors harbor microhomology^8,13,14^, indicative of potential DSB repair by alt-EJ. To determine whether complex rearrangements were present in alt-EJ-deficient cells lacking Polθ, we performed WGS on four *POLQ* KO sub-clones that harbored apparent complex rearrangements of the Y chromosome following sustained centromere inactivation, as determined using the previously described two-colored FISH approach (**Figure 2c**). Indeed, WGS revealed that three out of four *POLQ* KO sub-clones exhibited the oscillating DNA copy number patterns (**Figure 2d**) that are characteristic of cancer-associated chromothripsis^8,15^. One sub-clone harbored a region of clustered C>T hypermutation (**Figure 2d**), which has been previously shown to arise near chromothriptic breakpoints^16,35^. Thus, Polθ-mediated alt-EJ is largely dispensable for the formation of complex rearrangements following chromothripsis from micronuclei.

We next focused on DNA-PKcs-deficient cells for further studies. DNA-PKcs promotes the synapsis of two DSB ends by interacting with Ku and activating its kinase activity, which is essential for its function in NHEJ^36–38^. Complementation of *PRKDC* KO cells with WT DNA-PKcs, but not a K3752R kinase-dead (KD) mutant^36^ (**Extended Data Figure 4a**), restored the formation of complex Y chromosome rearrangements (**Extended Data Figure 4b-e**), confirming that the deficiencies observed in *PRKDC* KO clones are due to the lack of DNA-PKcs kinase activity. Although rearrangements were markedly reduced in all NHEJ-deficient settings examined, we note that complex rearrangements remained present at a low yet detectable frequency in the absence of NHEJ (**Figure 1d**, **Figure 2b**).

To determine whether DSB repair pathways beyond NHEJ were responsible for the complex rearrangements observed in DNA-PKcs-deficient cells, we inhibited a second DSB repair pathway in *PRKDC* KO cells using pharmacological agents to inactivate HR by blocking the BRCA2-RAD51 interaction with CAM833^39^ or alt-EJ with the Polθ inhibitor ART558^40^, as well as the PARP inhibitor olaparib^41^. Cells lacking DNA-PKcs treated with HR, alt-EJ, and PARP inhibitors formed complex rearrangements at a frequency similar to vehicle-treated controls (**Extended Data Figure 5a-b**). To extend these findings genetically, we used CRISPR/Cas9 editing to generate cells deficient in both DNA-PKcs and a second DSB repair gene (*POLQ*, *RAD52*, or *RAD54L*; **Extended Data Figure 5c**). In all double KO settings examined, complex rearrangements remained detectable at a low frequency comparable to loss of DNA-PKcs alone (**Extended Data Figure 5d**), suggesting that DSB repair pathways beyond NHEJ are not responsible for the complex rearrangements observed in the absence of DNA-PKcs.

DNA-PKcs promotes DNA end ligation by forming a long-range synaptic complex during NHEJ. However, *de novo* short-range synaptic complex containing XLF can form in the absence of DNA-PKcs, indicating that DNA end ligation may be possible without DNA-PKcs^42^. This is further suggested by a functional redundancy between DNA-PKcs and XLF in NHEJ^43,44^. To determine whether the residual rearrangements in DNA-PKcs KO cells are formed through minimal NHEJ activity through XLF-mediated end synapsis, we tested for a synergistic reduction in rearrangement formation following depletion of XLF in both WT and *PRKDC* KO cells. Similar to *NHEJ1* KO cells, depletion of XLF in WT cells was sufficient to reduce complex rearrangements (**Extended Data Figure 5e-g**). Importantly, complex rearrangements were exceedingly rare in *PRKDC* KO cells depleted of XLF, occurring in only 3 out of 168 (1.8%) Y chromosomes with rearrangements (**Extended Data Figure 5g**). Altogether, these data highlight that DSB repair pathways beyond NHEJ minimally contribute to chromothripsis-induced complex rearrangements.

### Repair kinetics of fragmented chromosomes in MN bodies

Chromosomes encapsulated in micronuclei acquire DSBs, which can be detected by the phosphorylation of histone H2AX (γH2AX). Micronuclei are dysfunctional in sensing and/or repairing DNA damage following rupture of its nuclear envelope^1^, suggesting that micronuclear DSBs cannot be repaired until its reincorporation into a functional nucleus. Consistent with this, immunofluorescent staining revealed that micronuclear envelope rupture – as determined by the loss of nucleocytoplasmic compartmentalization^1^ or recruitment of the cytosolic DNA sensor cGAS^45,46^ – triggered the loss of DNA-PKcs specifically from micronuclei (**Extended Dat Figure 6a-d**). *PRKDC* KO cells displayed similar levels of γH2AX within micronuclei as compared to WT controls, further supporting that DSBs are not actively repaired within ruptured micronuclei during interphase (**Extended Data Figure 6e-f**).

Since NHEJ is suppressed in mitosis^47^, we hypothesized that most fragments are likely carried over to one or both resulting daughter cell(s) for repair during the subsequent G1 phase. To directly test this, we analyzed DNA damage by immunofluorescence and DNA FISH (IF-FISH) on the previously micronucleated chromosome after its reincorporation into the primary nucleus as MN bodies. Compared to WT cells, increased γH2AX and 53BP1 signals were observed on FISH-labeled Y chromosomes manifesting as MN bodies in *PRKDC* KO cells, indicating that NHEJ engages reincorporated chromosome fragments in the nucleus for repair (**Figure 3a-d**).

**Figure 3.**
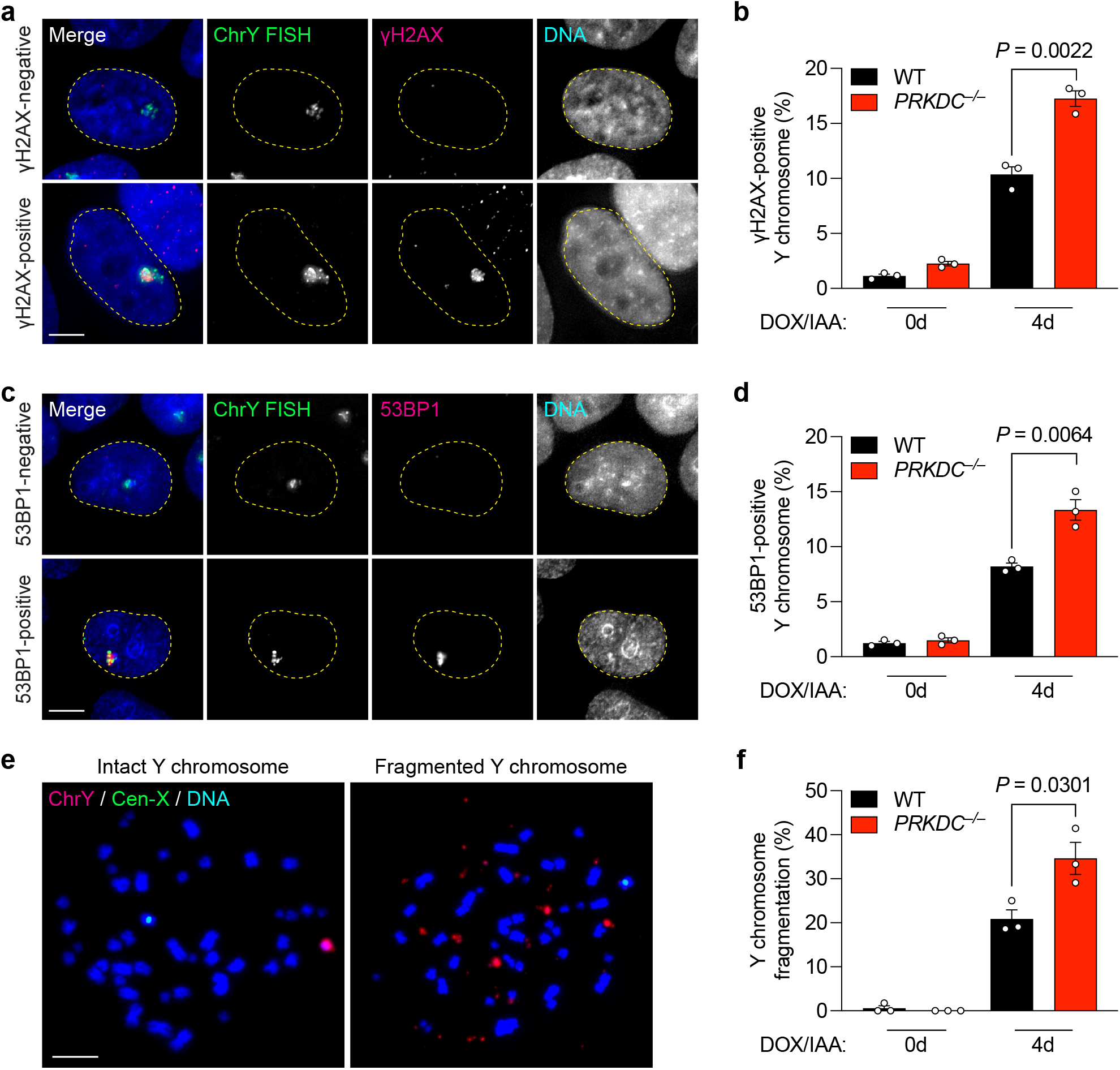
NHEJ-deficient cells accumulate damaged chromosome fragments within nuclear MN bodies. **a)** Images of interphase cells with γH2AX-negative Y chromosome or γH2AX-positive Y chromosomes within an MN body after 4d DOX/IAA treatment. Scale bar, 5 µm. **b)** Frequency of γH2AX-positive Y chromosomes. Data were pooled from (left to right): 549, 353, 429, and 293 cells. **c)** Images of interphase cells with 53BP1-negative or 53BP1-positive Y chromosomes after 4-day DOX/IAA treatment. Scale bar, 5 µm. **d)** Frequency of 53BP1-positive Y chromosomes. Data were pooled from (left to right): 567, 543, 634, and 592 cells. **e)** Images of metaphase spreads with an intact or fragmented Y chromosome after 4d DOX/IAA treatment. Scale bar, 10 µm. **f)** Frequency of Y chromosome fragmentation. Data pooled from (left to right): 329, 291, 347, and 373 metaphase spreads. Bar graphs in (**b**), (**d**), and (**f**) represent the mean ± SEM of *n* = 3 independent experiments. Statistical analyses were calculated by unpaired Student’s t-test.

Examination of metaphase spreads for chromosome fragmentation in WT and *PRKDC* KO cells over multiple cell cycles revealed that *PRKDC* deficiency resulted in an accumulation of Y chromosome fragments over time (**Figure 3e-f**). Thus, in the absence of NHEJ, fragmented chromosomes reincorporate into the nucleus and persist unrepaired throughout the cell cycle.

We next used live-cell imaging to monitor the kinetics of DSB repair by fusing the minimal focus-forming region of 53BP1^48^ to a HaloTag (Halo-53BP1). To label and track the Y chromosome from micronuclei into daughter cell nuclei, we used a recently developed dCas9-based SunTag reporter targeting a large repetitive array on the Y chromosome^9^. In the example shown in **Figure 4a**, a mother cell with a Y chromosome-specific micronucleus underwent mitosis and subsequently formed a large, Halo-53BP1-labeled MN body in the nucleus of one of the daughter cells. This MN body co-localized with SunTag-labeled Y chromosome fragments, demonstrating that it indeed originated from the micronucleated chromosome from the preceding cell cycle that had now re-incorporated into the primary nucleus (**Figure 4a**). Whereas the fluorescence intensity of the dCas9-SunTag reporter remained constant throughout the cell cycle, the Halo-53BP1 signal accumulated during early G1 phase, reached a plateau approximately 10 hours after mitosis, and proceeded to gradually decline over a 20-hour window during interphase (**Figure 4b-c**).

**Figure 4.**
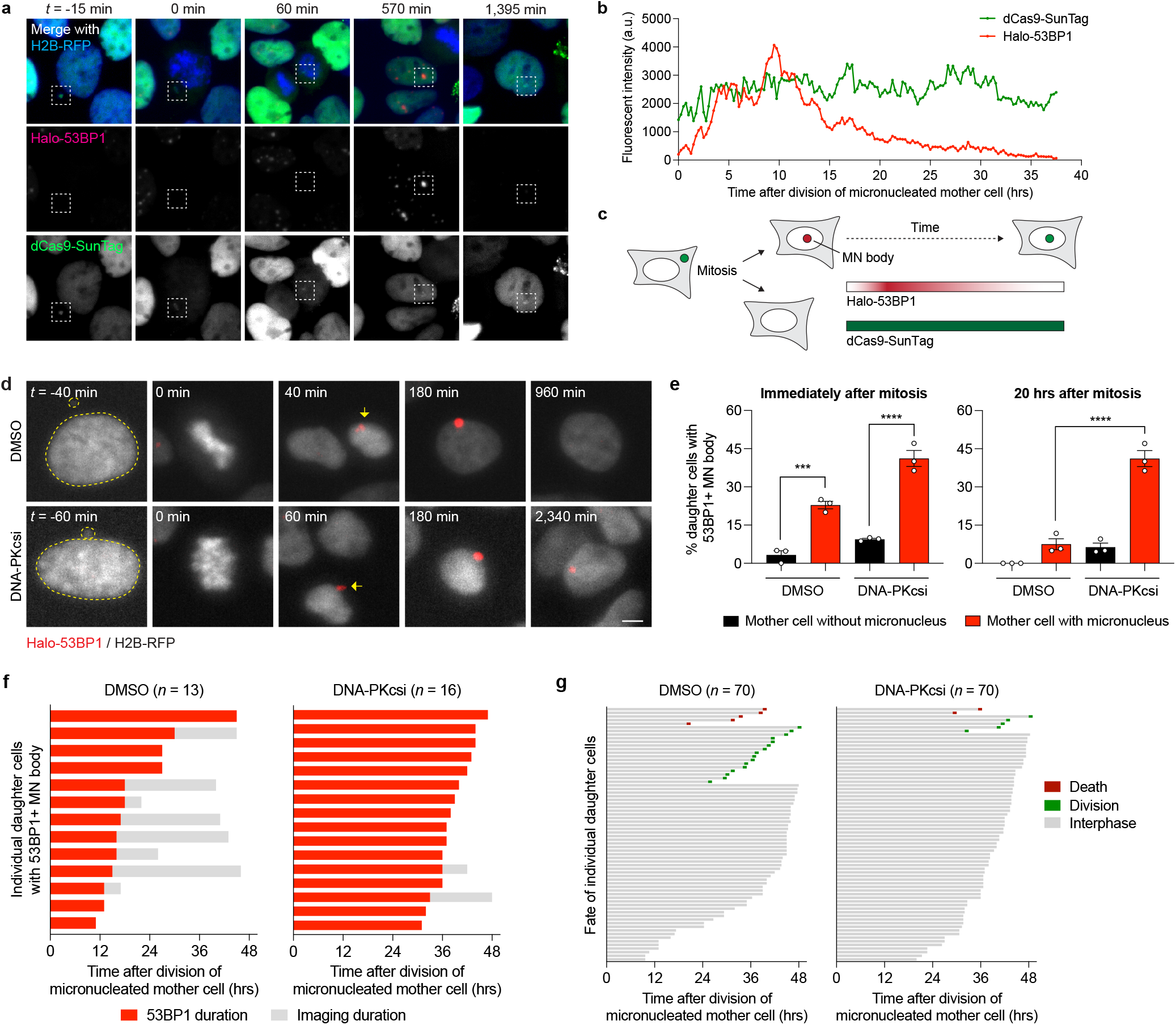
Inhibition of NHEJ prolongs 53BP1 residence time at MN bodies and triggers cell cycle arrest. **a)** Example time-lapse images of a mother cell harboring a dCas9-SunTag-labeled Y chromosome in a micronucleus undergoing cell division, which in turn generates a daughter cell that incorporates the micronucleated chromosome into the nucleus as an MN body labeled with a HaloTag fused to the minimal focus-forming region of 53BP1 (Halo-53BP1). Time point at zero min. depicts mitotic entry. **b)** Measurement of Halo-53BP1 fluorescence intensity over a ∼37-hour imaging period. The dCas9-SunTag-labeled Y chromosome is monitored as a control. **c)** Schematic of Halo-53BP1 residence time at MN bodies from panels (**a-b**) after mitosis. **d)** Representative time-lapse images of newly-formed, Halo-53BP1-labeled MN bodies with or without treatment with the DNA-PKcs inhibitor AZD7648. Scale bar, 5 µm. **e)** Frequency of Halo-53BP1-labeled MN body formation and persistence. Data represent mean ± SEM of *n* = 3 independent experiments pooled from (left to right): 63, 64, 72, and 74 micronucleated mother cells. Statistical analyses were calculated by ordinary one-way ANOVA test with multiple comparisons. ns, not significant; \*\*\**P* ≤ 0.001; \*\*\*\**P* ≤ 0.0001. **f)** Duration of Halo-53BP1 residence time from panel (**d**) after MN body formation with or without inhibition of DNA-PKcs in individual daughter cells. **g)** Fate of daughter cells after division of micronucleated mother cells. Sample sizes in panels (**f-g**) represent individual daughter cells.

In WT cells, ∼23% of micronucleated mother cells formed daughter cells with 53BP1-labeled MN bodies compared to ∼3% from non-micronucleated control cells (**Figure 4d-e**), confirming that MN bodies are indeed derived from the micronucleated chromosome. In DMSO-treated control cells, Halo-53BP1 persisted for an average of ∼17.9 hours until its resolution (**Figure 4f**). In the presence of the DNA-PK inhibitor AZD7648^49^, MN body-associated Halo-53BP1 signals persisted during the entire duration of imaging, often exceeding 30-40 hours (**Figure 4f**). We next tracked the fate of daughter cells from micronucleated mother cells. Long-term live-cell imaging revealed that inhibition of DNA-PKcs reduced the proportion of cells that successfully entered mitosis, indicative of cell cycle arrest (**Figure 4g**). These data suggest that the sustainment of DNA damage signaling as persistent MN bodies can activate the cell cycle checkpoint.

DNA damage on micronucleated chromosomes are repaired mainly through NHEJ, leading to the formation of rearrangements that can be detected by FISH. We analyzed γH2AX levels on FISH-labeled fragmented chromosomes that had reincorporated into the nucleus at different time points after mitosis (**Figure 5a**). In agreement with live-cell imaging experiments (**Figure 4**), most fragmented chromosomes were repaired within 20 hours after mitosis in WT cells, whereas *PRKDC* KO cells continued to harbor γH2AX marks within MN bodies that persisted beyond 20 hours (**Figure 5b-c**). Since rearrangements are a byproduct of successful DSB repair, we next sought to directly visualize rearrangements of the Y chromosome in interphase cells at similar time points. To do so, we induced premature chromosome condensation by treatment with the phosphatase inhibitor calyculin A and analyzed metaphase-like chromosomes harboring either unduplicated chromatids from G1-phase cells or sister chromatids from G2-phase cells (**Figure 5d**). In both WT and *PRKDC* KO cells, most Y chromosomes remained fragmented six hours after mitosis during G1 phase (**Figure 5e**). As WT cells were allowed to progress throughout the cell cycle into G2 phase, such fragmented chromosomes underwent successful yet error-prone repair to form rearranged chromosomes within 20 hours. In contrast, the Y chromosome in *PRKDC* KO cells remained fragmented, indicative of defects in forming rearrangements in the absence of NHEJ (**Figure 5e**). These data provide direct evidence supporting the ligation of reintegrated fragments within MN bodies during a single cell cycle by the NHEJ pathway.

**Figure 5.**
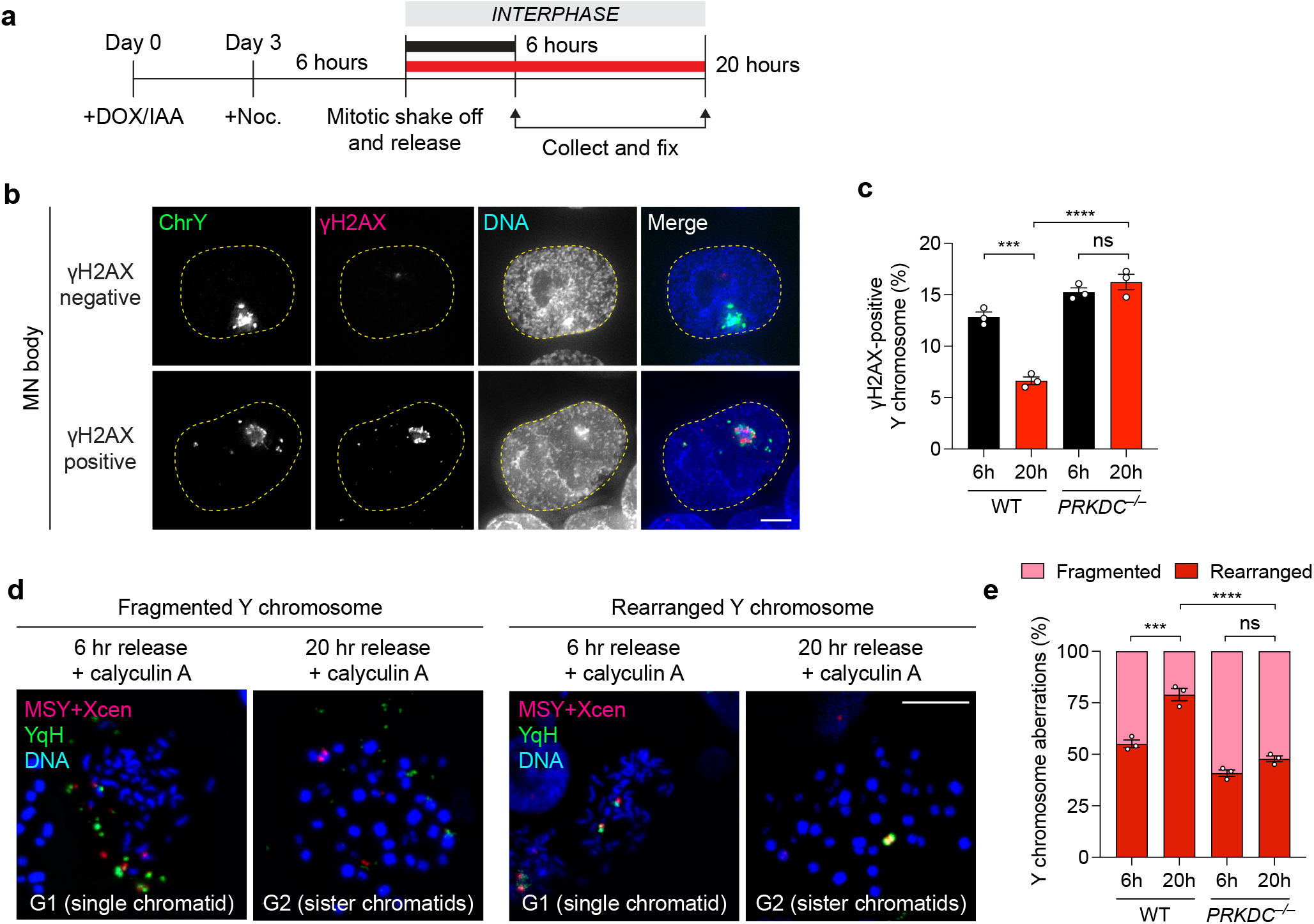
Reincorporated chromosome fragments undergo NHEJ-dependent rearrangement during the subsequent interphase. **a)** Schematic of experiment to assess DSB repair at 6 and 20 hours after mitosis. Noc, nocodazole. **b)** Images of interphase cells with γH2AX-negative or γH2AX-positive Y chromosomes before and after division of micronucleated mother cells by IF-FISH. Scale bar, 5 µm. **c)** Frequency of γH2AX-positive Y chromosomes in WT and *PRKDC* KO clone 1 by IF-FISH. Data pooled from (left to right): 352, 361, 341, and 327 Y chromosome-positive interphase cells. **d)** Representative images of asynchronous cells treated with calyculin A to induce premature chromosome condensation. Chromosome spreads were subjected to DNA FISH using the indicated probes. Scale bar, **10** µm. **e)** Frequency of Y chromosome aberrations in WT controls and *PRKDC* KO clone from panel (**d**). Data pooled from (left to right): 241, 128, 178, and 159 chromosome spreads. Bar graphs in (**c**) and (**e**) represent mean ± SEM of *n* = 3 independent experiments. Statistical analyses were calculated by ordinary one-way ANOVA test with multiple comparisons. ns, not significant; \*\*\**P* ≤ 0.001; \*\*\*\**P* ≤ 0.0001.

Lastly, we sought to investigate the basis of how inter-chromosomal rearrangements are formed between fragmented chromosomes in MN bodies with seemingly intact non-homologous chromosomes in the nucleus, which may arise from the improper repair of micronucleated fragments to spontaneous DSBs in the genome. To test this hypothesis, we induced Y chromosome-MN body formation and exposed cells to either low dose ionizing radiation or a telomere-incorporating nucleoside analogue (6-thio-dG)^50^ to induce DNA damage genome-wide or specifically at telomeres, respectively. Both sources of DNA damage were sufficient to elevate the frequency of inter-chromosomal rearrangements involving the Y chromosome (**Extended Data Figure 7a-b**). To determine whether chromosome fragments from micronuclei can integrate into a targeted genomic locus, we introduced a site-specific DSB by using Cas9 RNPs to trigger the cleavage of two autosomes (chromosomes 3 and 5) or the X chromosome during a window in which MN bodies were present. Indeed, site-specific DSB induction promoted Y chromosome translocations and fusions with the targeted chromosome (**Extended Data Figure 7c-d**), indicating that inter-chromosomal rearrangements can be generated by the ligation of chromosome fragments from MN bodies to sites of concurrent DNA damage.

## DISCUSSION

Complex rearrangements arising from chromothripsis are characterized by extensive rearrangements that are confined to one or a few chromosome(s)^8^. Several mutagenic DSB repair pathways can potentially reassemble the fragmented chromosome in seemingly random orientation. Here we demonstrated that fragmented micronuclear chromosomes are predominantly repaired by NHEJ to form complex rearrangements reminiscent of those found in human cancers. Damaged fragments from micronuclei persist throughout mitosis and are carried over into the subsequent cell cycle following its reincorporation into daughter cell nuclei as MN bodies.

Following nuclear reincorporation, analysis of the kinetics of repair suggest that these fragments become reassembled during a prolonged interphase within MN bodies, perhaps due to activation of DNA damage checkpoints and/or the presence of a substantial number of DSB ends that require processing prior to ligation. Live-cell imaging revealed an unexpected delay (by approximately 10 hours) in the onset of 53BP1 recruitment to MN bodies and its subsequent resolution (**Figure 4**), consistent with a prolonged residence time for MDC1 that has been observed on MN bodies^11^. These delays in DSB repair may be caused by the binding of the CIP2A-TOPBP1 complex to the fragmented chromosome during the previous interphase or mitosis^9,10^ and that are carried over into early G1 phase of the cell cycle. Alternatively, ssDNA on micronucleated chromosomes that are acquired during interphase^51^ or mitotic entry^52^ may require processing upon reincorporation into a functional nucleus to generate fragmented DSB ends that are compatible for ligation by NHEJ.

Analysis of breakpoint junctions from cancer genome sequencing data revealed that most rearrangement events occur without significant microhomology, indicative of ligation between blunt-ended DSB ends. Some junctions exhibit short tracts of microhomology, suggesting a potential contribution by alt-EJ and/or microhomology-mediated break-induced replication^5,13,53,54^. However, our data highlight NHEJ as the predominant and perhaps exclusive DSB repair pathway for chromothripsis arising from mitotic errors. Previous studies demonstrated that chromosome fragments from micronuclei can be recognized by the DNA damage response throughout mitosis^9–11^, a period in which Polθ-mediated alt-EJ is active^55–57^. Although we cannot fully exclude the possibility that a small fraction of fragmented chromosome ends may be repaired by alt-EJ during mitosis, inactivation of the alt-EJ pathway through biallelic deletions of *POLQ* or with small molecule inhibitors targeting Polθ had an undetectable impact on both the frequency and spectrum of rearrangements produced by micronucleation. We speculate that the binding of the CIP2A-TOPBP1 complex to fragmented chromosomes in micronuclei during the interphase-to-mitosis transition^9^ may preclude engagement by Polθ and/or other alt-EJ components involved in mitosis-specific DSB repair. Indeed, inhibition of the CIP2A-TOPBP1 pathway abolished the formation of MN bodies in interphase cells, which in turn suppressed the formation of complex rearrangements^9^.

In the absence of canonical NHEJ factors, the rearrangements that are generated from micronuclei formation lack the features of complex rearrangements that are characteristic of chromothripsis, including oscillating DNA copy-number alterations. The rearrangement landscape shifts to favor more simple alterations that are typically comprised of unbalanced translocations, whole-chromosome fusions, or chromosome-arm deletions, which could arise from a fraction of micronuclei that harbor relatively few DNA breaks^7^ and/or mis-segregated chromosomes that undergo breakage during cytokinesis^58^. These larger chromosome fragments can then undergo ligation to spontaneous DSBs in the genome to generate cytogenetically-visible inter-chromosomal rearrangements. By contrast, NHEJ-deficient cells harboring more extensive DNA damage from catastrophically shattered chromosomes that cannot be repaired will ultimately undergo cell cycle arrest. Pharmacological inhibition of NHEJ (e.g., with small molecule inhibitors against DNA-PKcs) may therefore represent a therapeutic avenue to combat chromosomally unstable tumors or those treated with microtubule inhibitors to induce severe mitotic defects. Similar strategies targeting DNA-PKcs may also be effective in suppressing linear chromosome fragments from ligating into a circular extrachromosomal DNAs that can amplify oncogenes and/or genes conferring resistance to anti-cancer therapies^59,60^.

## AUTHOR CONTRIBUTIONS

Q.H. and P.L. conceived the project and designed the experiments. Q.H., R.D., A.G., A.M., E.G.M., and J.L.E. performed experiments and analyzed the data. J.E.V.-I. and I.C.-C. analyzed whole-genome sequencing data. I.C.-C. and P.L. provided supervision. Q.H. and P.L. wrote the manuscript with input from all authors.

## STATEMENT OF COMPETING INTERESTS

All authors have no competing interests to declare.

## DATA AVAILABILITY STATEMENT

Whole-genome sequencing data presented in this manuscript have been deposited at the European Nucleotide Archive under project accession ID PRJEB64431.

## MATERIALS AND METHODS

### Cell lines and reagents

DLD-1 cells were cultured in Dulbecco’s Modified Eagle Medium (Thermo Fisher) supplemented with 10% tetracycline-free fetal bovine serum (Omega Scientific) and 100 U/ml penicillin-streptomycin at 37°C under 5% CO_2_ atmosphere. Cells were routinely confirmed free of mycoplasma contamination. The derivation of DLD-1 cells expressing the dCas9-SunTag system, mCherry-NLS, and cGAS-GFP were previously described^9^. To generate the 53BP1 reporter system, a HaloTag was fused in-frame to the N-terminus of the minimal focus-forming region (FFR, amino acids 1220-1711) of 53BP1 from *TP53BP1* cDNA (a gift from Anthony Davis) and cloned into a pBABE-zeo construct (Addgene). DLD-1 cells engineered to carry the dCas9-SunTag system and expressing H2B-mCherry were transduced with retroviruses that were packaged in 293GP cells for 24 hours and selected with 50 μg/mL zeocin for two weeks. Single cell derived-clones forming robust 53BP1 foci were isolated and used for live-cell imaging experiments.

Doxycycline (DOX) and auxin (indole-3-acetic acid, IAA) were used at 1 μg/ml and 500 μM, respectively. Nocodazole (Millipore-Sigma) was used at 100 ng/mL for mitotic arrest. Geneticin (G418 Sulfate) was used at 300 mg/mL for selection. Small molecules compounds were used at the following concentrations: 10 μM CAM833 (Tocris Bioscience), 0.5 μM Olaparib (Cayman Chemical), 1 μM ART558 (MedChemExpress), 1 μM AZD7648 (MedChemExpress) and 0.5 μM 6-thio-2’-deoxyguanosine (6-thio-dG, a gift from Jerry Shay, UTSW). Dose-response assays were performed to identify an optimal and tolerable drug concentration without affecting DLD-1 cell growth and viability. Ionizing radiation experiments were performed using a Mark I Cesium-137 irradiator (JL Shepherd).

### Genome editing

To generate KO clones, TrueCut Cas9 v2 (Thermo Fisher) and sgRNAs (synthesized by Synthego) were assembled into ribonucleoprotein complexes and transfected into cells using Lipofectamine CRISPRMAX Cas9 Transfection Reagent (Thermo FIsher). 72 hours post-transfection, cells were plated at low density (50 cells per 10-cm^2^ dish) to isolate single cell-derived clones. After approximately two weeks, colonies were isolated using cloning cylinders and expanded. Clones were screened by PCR for targeted deletions and confirmed to harbor frameshift mutations by Sanger sequencing. When antibodies were available, immunoblotting was used to confirm the loss of the target protein. All sgRNA sequences and PCR primers used in this study are provided in **Supplementary Table 1**.

### Cell growth assays

For viability assays, 3×10^4^ cells per well were seeded into 6-well plates with or without DOX/IAA treatment. Three days later, cells were washed three times with PBS and supplemented with fresh media without DOX/IAA. Cells were transferred to 10-cm^2^ plates three days later and selected with G418 for 10 days. To calculate relative viability in G418, the total number of cells in the DOX/IAA condition was divided by the total number of cells in the control condition. For quantification of long-term cell growth rates, cells were continuously cultured for approximately one month and the total cell numbers were counted during each passage.

### RNA interference and complementation

Cells were transfected with 20 nM small interfering RNA (siRNA, Thermo Fisher) using Lipofectamine RNAiMAX (Thermo Fisher) according to the manufacturer’s instructions. All siRNA sequences used in this study are provided in **Supplementary Table 1**. For complementation experiments, a vector containing FLAG-tagged WT or kinase-dead *PRKDC* (K3752R) cDNA (a gift from Anthony Davis, UTSW) was co-transfected with pmaxGFP using a Nucleofector II (Amaxa). Ten days post-transfection, GFP-positive cells were sorted by flow cytometry using a FACSAria (BD Biosciences) into individual wells of a 96-well plate. Clones were expanded and screened by immunoblotting for expression of FLAG-tagged DNA-PKcs.

### Immunoblotting

Cells were collected by trypsinization and pelleted by centrifugation. Cell pellets were washed once with ice-cold PBS and lysed in 2x Laemmli Sample Buffer (50mM Tris-HCl PH6.8, 2% SDS, 10% glycerol, 0.01% bromophenol blue, 2.5% β-mercaptoethanol). Samples were denatured by boiling at 100°C for five min and resolved by SDS–PAGE. The proteins were transferred to polyvinylidene difluoride membranes using a Trans-Blot Turbo System (Bio-Rad). Blots were blocked with 5% milk in PBST (PBS, 0.1% Tween-20) for one hour at room temperature before incubation with primary antibodies (1:1,000 dilution in PBST except for anti-α-tubulin, which was used at 1:5,000) overnight at 4°C. Blots were washed three times in PBST with 10 mins each, followed by incubation with horseradish peroxidase-conjugated secondary antibodies (Sigma, 1:5,000 dilution in 5% milk in PBST) and an additional three washes in PBST. After adding chemiluminescent substrate (SuperSignal West Pico PLUS, Thermo Fisher), blots were visualized using a ChemiDoc Imaging System (Bio-Rad). A list of all primary antibodies used in this study is provided in **Supplementary Table 2**.

### Immunofluorescence

Cells were seeded on chamber slides or coverslips and fixed with 4% formaldehyde for 10 min at room temperature or with ice-cold methanol for 10 min at −20°C, washed three times with PBS, and permeabilized with 0.3% Triton X-100 in PBS for five min. After washing with PBS, cells were incubated with Triton Block (0.1% Triton X-100, 2.5% FBS, 0.2 M glycine, PBS) for one hour at room temperature, washed with PBS, and incubated with primary antibodies (1:1000 diluted in Triton Block) overnight at 4°C, followed by three washes with PBST-X (0.1% Triton X-100 in PBS) 10 min each. After washing, Alexa Fluor-conjugated secondary antibodies (Invitrogen) were diluted 1:1000 in Triton Block and applied to cells for one hour at room temperature, followed by three washes with PBST-X. Cells were stained with DAPI, rinsed with PBS, air dried, and mounted with ProLong Gold antifade mounting solution (Invitrogen) before imaging.

### Metaphase spread preparation

To prepare metaphase spreads, cells were treated with 100 ng/ml colcemid (KaryoMAX, Thermo Fisher) for four hours, collected by trypsinization, and centrifuged at 180 RCF for 5 min. Cell pellets were gently resuspended in 500 μL PBS, and 5 mL pre-warmed 0.075 M KCl was added dropwise to the tube while vortexing at low speed. Cells were then incubated at 37°C for six min followed by adding 1 mL of freshly-made Carnoy’s fixative (3 methanol:1 acetic acid), followed by centrifugation at 180 RCF for five min and removal of the supernatant. Cell pellets were resuspended in 6 mL ice-cold Carnoy’s fixative, followed by centrifugation at 180 RCF for five min and resuspension in 500 μL Carnoy’s fixative. Fixed samples were dropped onto slides and air dried.

To induce premature chromosome condensation, 100 nM calyculin A (Cell Signaling) were added to directly to the cell culture medium and incubated for one hour at 37°C. Cells were then harvested and centrifuged at 180 RCF for 5 min. The cell pellets were incubated in 0.075 M KCl followed by fixation in Carnoy’s fixative, as described above.

### DNA fluorescence in situ hybridization (FISH)

FISH probes (MetaSystems) were applied to metaphase spreads dropped onto slides. Slides were sealed with a coverslip and denatured at 75°C for two mins. After denaturation, samples were incubated at 37°C overnight in a humidified chamber for hybridization. Samples were then washed with 0.4x SSC at 72°C for two min and 2x SSCT (2x SSC, 0.05% Tween-20) for 30 sec at room temperature. The samples were stained with DAPI and mounted with ProLong Gold antifade mounting solution.

For immunofluorescence combined with DNA FISH (IF-FISH), the immunofluorescence procedure was performed first as described above and fixed with Carnoy’s fixative for 15 min at room temperature. Samples were rinsed with 80% ethanol and air dried before proceeding to the FISH protocol. For rearrangement experiments, inhibitors were added to cells simultaneously with DOX/IAA and incubated for 6 days prior to G418 selection.

### Live-cell imaging

To perform live-cell imaging, cells were seeded in 96-well glass-bottom plates (Cellvis, P96- 1.5H-N). DLD-1 cells expressing H2B-mCherry, the dCas9-SunTag system, and 53BP1-Halo were treated with DOX/IAA for 72 hours, and 200 nM JF646 ligand (Promega) was added 15 min prior to imaging. Images were captured every 20 minutes for 48 hours using an ImageXpress Confocal HT.ai High-Content Imaging System (Molecular Devices) equipped with a 40x objective in CO2-independent medium (Thermo Fisher) at 37°C. Images were acquired at 7 x 1.5 μm z-sections under low power exposure. Maximum intensity projections were generated using MetaXpress and movies were analyzed using Fiji (v.2.1.0/1.53c).

### Fluorescence microscopy

Metaphase FISH images were obtained using the Metafer Scanning and Imaging Platform (MetaSystems). Briefly, slides were pre-scanned for metaphases using the M-search Mode with a 10x objective (ZEISS Plan-Apochromat 10x/0.45). Image capturing was performed using the Auto-cap Mode with a 63x objective (ZEISS Plan-Apochromat 63x/1.40 oil). Image analyses were performed using Isis Fluorescence Imaging Platform (MetaSystems) and Fiji (v.2.1.0/1.53c).

Immunofluorescent or IF-FISH images were acquired using the DeltaVision Ultra microscope system (GE Healthcare), which was equipped with a 4.2 MPx sCMOS detector. Images were captured using a 100x objective (UPlanSApo, 1.4 NA) with 15 × 0.2-μm z-sections. Images were deconvolved and maximum intensity projections were generated using the softWoRx program (v.7.2.1, Cytiva). Images were analyzed using Fiji (v.2.1.0/1.53c).

### Whole-genome sequencing

For whole-genome sequencing, genomic DNA was extracted from ∼3×10^6^ cells by using Quick-DNA Kit (Zymo Research) according to manufacturer’s instructions. Sequencing library preparation and whole genome sequencing were performed by Novogene. Briefly, the genomic DNA of each sample was sheared into short fragments of about 350 bp and ligated with adapters. Whole genome sequencing was performed by using NovaSeq PE150 platform at ∼30x coverage.

Whole-genome sequencing data were aligned to the GRCh38 build of the human reference genome using BWA-MEM (v0.7.17)^61^. Aligned sequencing reads were processed using SAMtools (v1.12)^62^, and duplicate reads were flagged using Sambamba (v0.8.1)^63^. Sequencing depth was calculated at 10,000 base pair windows using Mosdepth (v0.3.1)^64^. Control-FREEC (v11.6)^65^ was used to perform copy number variation analysis using default parameters. Somatic single-nucleotide mutations were detected using SAGE (v2.8.0, https://github.com/hartwigmedical/hmftools). Visualization of inter-mutation distances across the genome (rainfall plots) was performed using the MutationalPatterns Bioconductor package (v3.10)^66^.

### Statistical analysis

Statistical analyses were performed using GraphPad Prism 9.2.0 software using the tests described in the figure legends. *P*-values ≤ 0.05 were considered statistically significant.

## EXTENDED DATA FIGURE LEGENDS

**Extended Data Figure 1.**
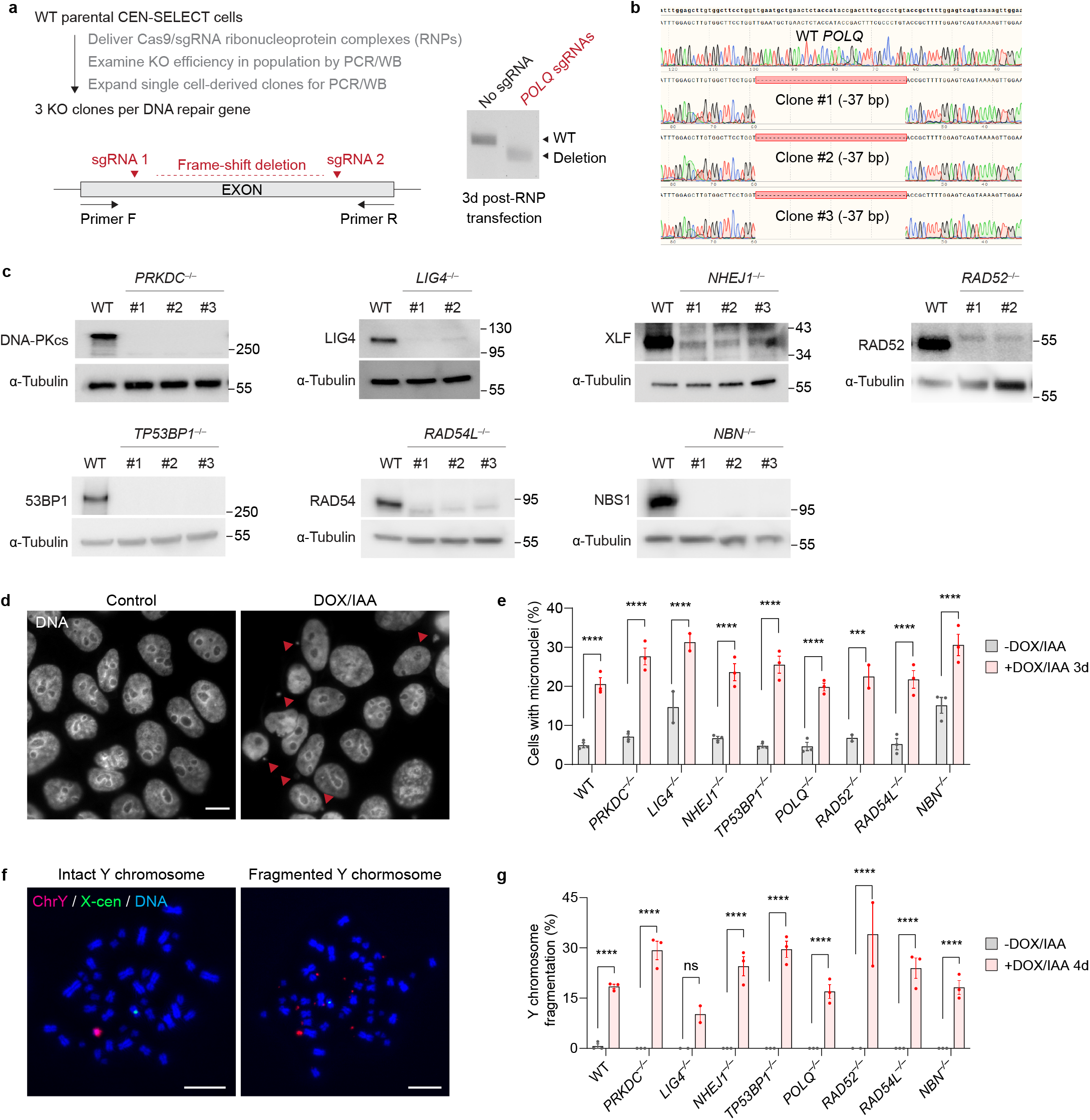
Generation of DSB repair knockout cells by CRISPR/Cas9 editing. **a)** Experimental schematic for generating CRISPR/Cas9-mediated biallelic knockout clones. Cleavage at two sgRNA sequences yields a frameshift deletion that can be detected by PCR. **b**) Sanger sequencing confirmation of predicted 37 base pair frameshift deletion in the *POLQ* gene in three independent clones. **c)** Confirmation of KO clones by immunoblotting. Molecular weight markers are indicated in kilodaltons. **d)** Representative images of *PRKDC* KO cells with micronuclei before and after induction with DOX/IAA. Scale bar, 10 µm. **e)** The percentage of cells with micronuclei with and without 3d DOX/IAA treatment. Data pooled from (left to right): 1,367, 1,711, 2,208, 1,236, 340, 407, 1,554, 1,709, 396, 335, 1,985, 1,769, 935, 295, 1,256, 1,782, 706, and 736 cells. **f)** Images of metaphase spreads with intact or fragmented Y chromosomes after 4d DOX/IAA treatment. Scale bar, 10 µm. **g)** Frequency of Y chromosome fragmentation. Only Y chromosome-positive metaphase spreads were scored. Data pooled from (left to right): 168, 267, 113, 226, 81, 141, 162, 162, 148, 147, 184, 213, 79, 123, 120, 186, 215, and 255 metaphase spreads. Bar graphs in (**e**) and (**g**) represent the mean ± SEM from *n* = 3 independent experiments for WT controls, *n* = 2 KO clones for *LIG4* and *RAD52*, and *n* = 3 KO clones for *PRKDC*, *NHEJ1*, *TP53BP1*, *POLQ*, *RAD54L*, and *NBN*; statistical analyses were calculated by ordinary one-way ANOVA test with multiple comparisons. ns, not significant; \*\*\**P* ≤ 0.001; \*\*\*\**P* ≤ 0.0001.

**Extended Data Figure 2.**
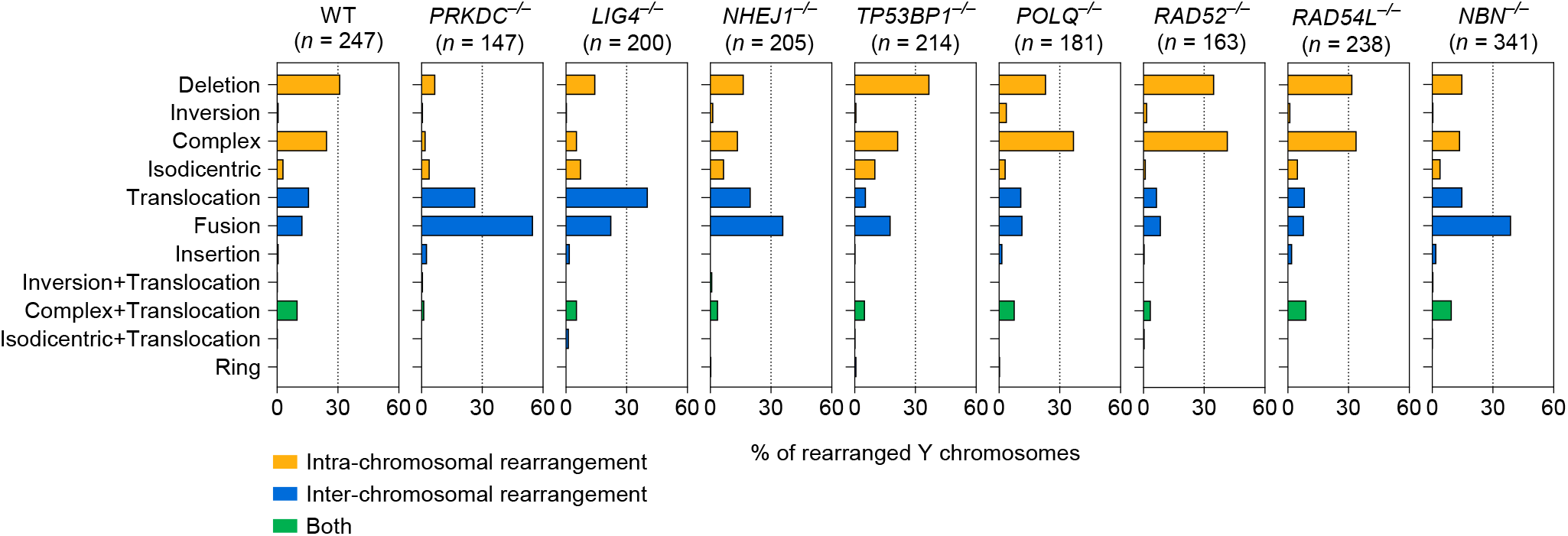
Genomic rearrangement landscape of mis-segregated chromosomes following transient centromere inactivation. Distribution of Y chromosome rearrangement types as determined by metaphase FISH following 3d DOX/IAA treatment and G418 selection. Sample sizes indicate the number of rearranged Y chromosomes examined; data are pooled from 3 independent experiments for WT cells and 2-3 independent KO clones for gene.

**Extended Data Figure 3.**
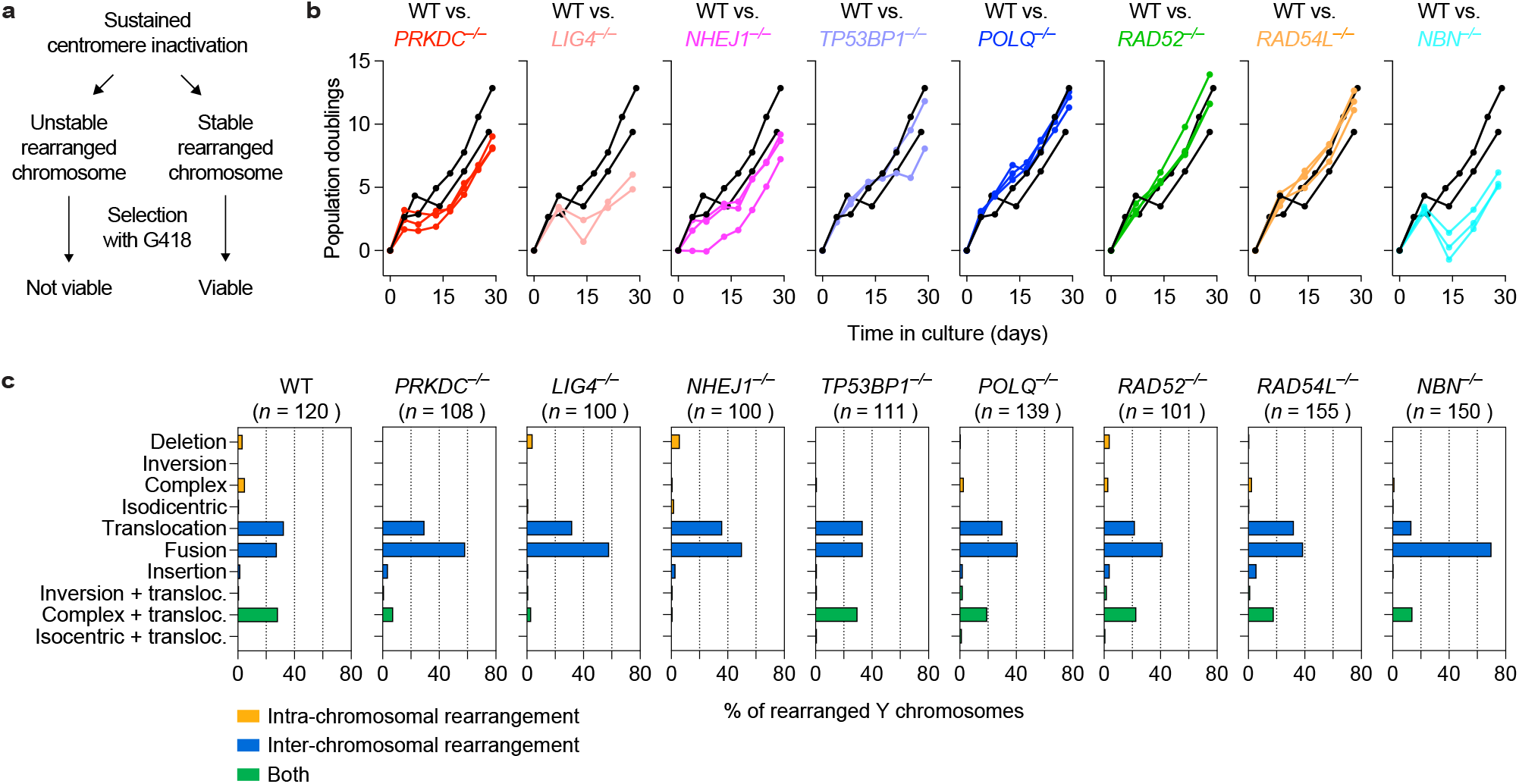
Genomic rearrangement landscape of mis-segregated chromosomes following sustained centromere inactivation. **a)** Schematic of turnover between proliferating cells harboring stable Y chromosome rearrangements and dying cells that are unable to maintain the Y-encoded *neoR* marker. **b)** Plots of cumulative cell doublings over a 30d period in the presence of DOX/IAA and G418. Each curve represents an individual clone except for WT, which represent independent experiments using the same cell line. **c)** Distribution of Y chromosome rearrangement types as determined by metaphase FISH following continuous passaging in DOX/IAA and G418. Sample sizes indicate the number of rearranged Y chromosomes examined; data are pooled from 2 independent experiments for WT cells or 2-3 independent KO clones for gene.

**Extended Data Figure 4.**
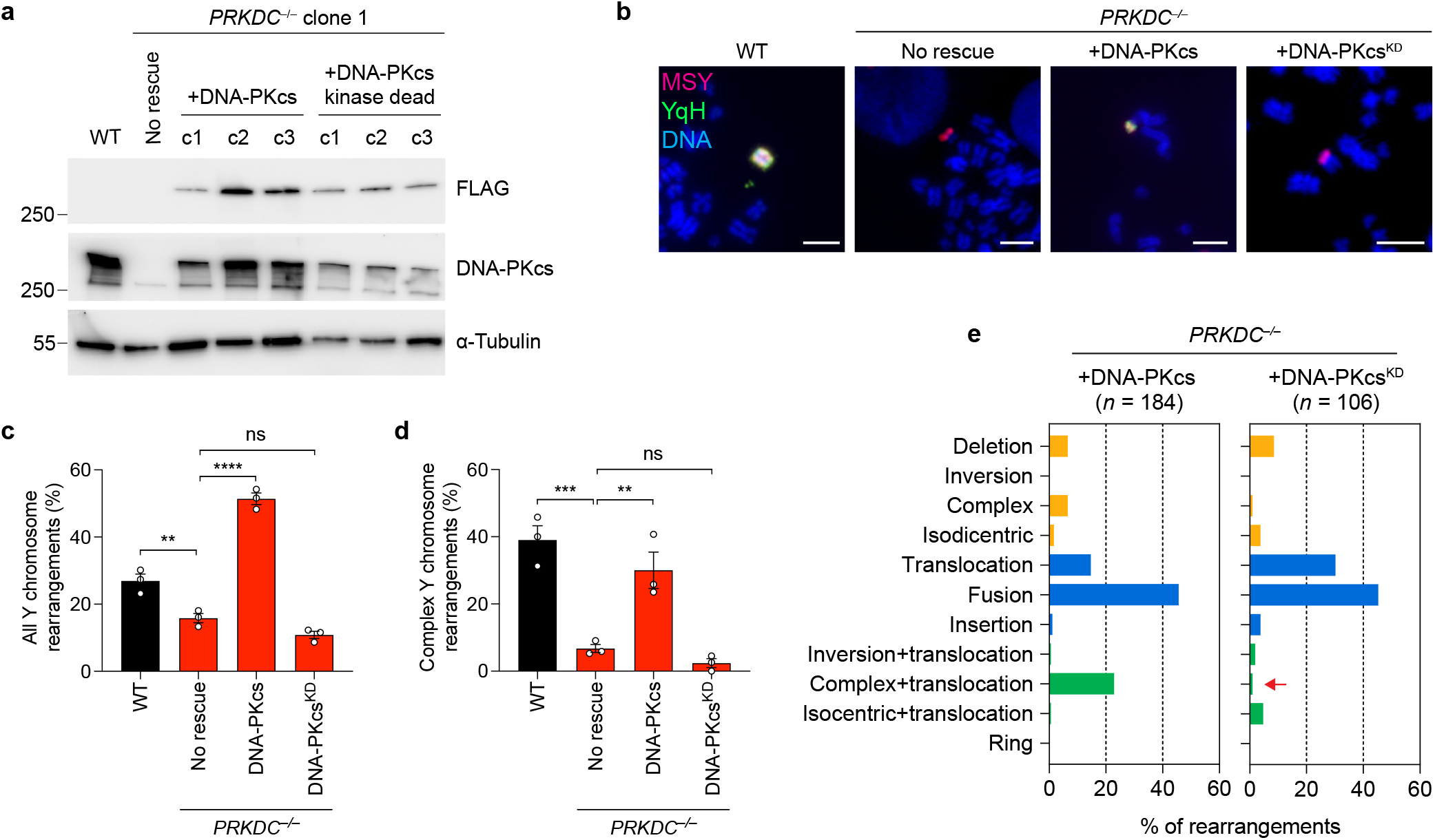
Characterization of genomic rearrangements in DNA-PKcs-deficient cells. **a)** Immunoblot confirmation of *PRKDC* KO cells expressing either FLAG-tagged WT (WT) or kinase-dead (KD) DNA-PKcs. Molecular weight markers are indicated in kilodaltons. **b)** Examples of Y chromosome rearrangements detected by metaphase FISH using the indicated sets of probes. Scale bar, 5 µm. **c)** Frequency of Y chromosome rearrangements in *PRKDC* KO cells complemented with WT or KD DNA-PKcs. Data are pooled from (left to right): 195, 339, 390, and 365 metaphase spreads. **d)** Proportion of Y chromosome rearrangements that can be classified as complex in *PRKDC* KO cells complemented with WT or KD DNA-PKcs. Data are pooled from (left to right): 70, 80, 184, and 106 metaphases spreads with Y chromosome rearrangements. Bar graphs in panels (**c-d**) represent the mean ± SEM of *n* = 3 independent experiments for WT and *PRKDC* KO, *n* = 3 independent DNA-PKcs rescue clones. Statistical analyses were calculated by ordinary one-way ANOVA test with multiple comparisons. ns, not significant; \*\**P* ≤ 0.001; \*\*\**P* ≤ 0.001; \*\*\*\**P* ≤ 0.0001. **e)** Distribution of Y chromosome rearrangement types as determined by metaphase FISH following 3d DOX/IAA treatment and G418 selection. Sample sizes indicate the number of rearranged Y chromosomes examined; data are pooled from 3 independent clones per condition.

**Extended Data Figure 5.**
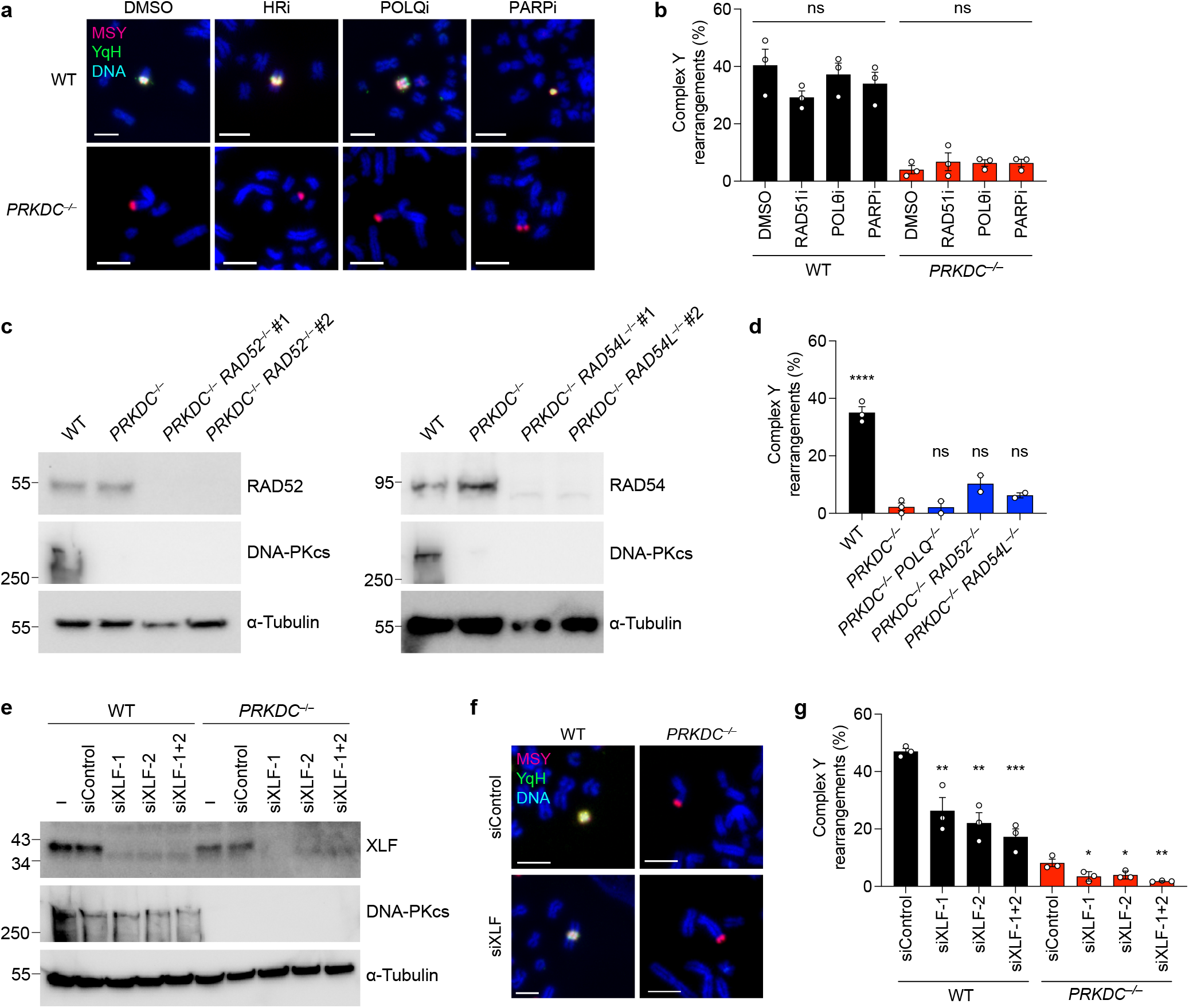
Complex rearrangements in DNA-PKcs-deficient cells arise from functional redundancies of the NHEJ pathway. **a)** Representative metaphase FISH images hybridized to the indicated probes from WT and *PRKDC* KO cells with Y chromosome rearrangements after different small inhibitor treatment. Scale bar, 5 µm. **b)** Frequency of complex rearrangements in WT and *PRKDC* KO clone 1 after treatment with the indicated small molecule inhibitors. Data represent mean ± SEM from *n* = 3 independent experiments pooled from (left to right): 167, 154, 158, 156, 170, 160, 157, and 159 metaphase spreads with Y chromosome rearrangements. **c)** Immunoblot confirmation of double KO clones. **d)** Frequency of complex rearrangements in double KO cells. Data represent mean ± SEM from *n* = 3 independent experiments for WT and *PRKDC* KO cells and *n* = 2 clones per double KO genotype analyzing (left to right): 177, 147, 128, 135, and 112 metaphase spreads containing Y chromosome rearrangements. **e)** Immunoblot confirmation of XLF depletion using the indicated siRNAs. **f)** Representative metaphase FISH images hybridized to the indicated probes from WT and *PRKDC* KO clone 1 with Y chromosome rearrangements after depletion of XLF. Scale bar, 5 µm. **g)** Frequency of complex rearrangements in WT and *PRKDC* KO clone #1 after depletion of XLF. Data represent mean ± SEM from *n* = 3 independent experiments pooled from (left to right): 134, 106, 95, 132, 152, 173, 180, and 168 metaphase spreads containing Y chromosome rearrangements. Statistical analyses for panels (**b**), (**d**), and (**g**) were calculated by ordinary one-way ANOVA test with multiple comparisons. ns, not significant; \**P* ≤ 0.05; \*\**P* ≤ 0.01; \*\*\**P* ≤ 0.001; \*\*\*\**P* ≤ 0.0001. Molecular weight markers in panels (**c-e**) are indicated in kilodaltons.

**Extended Data Figure 6.**
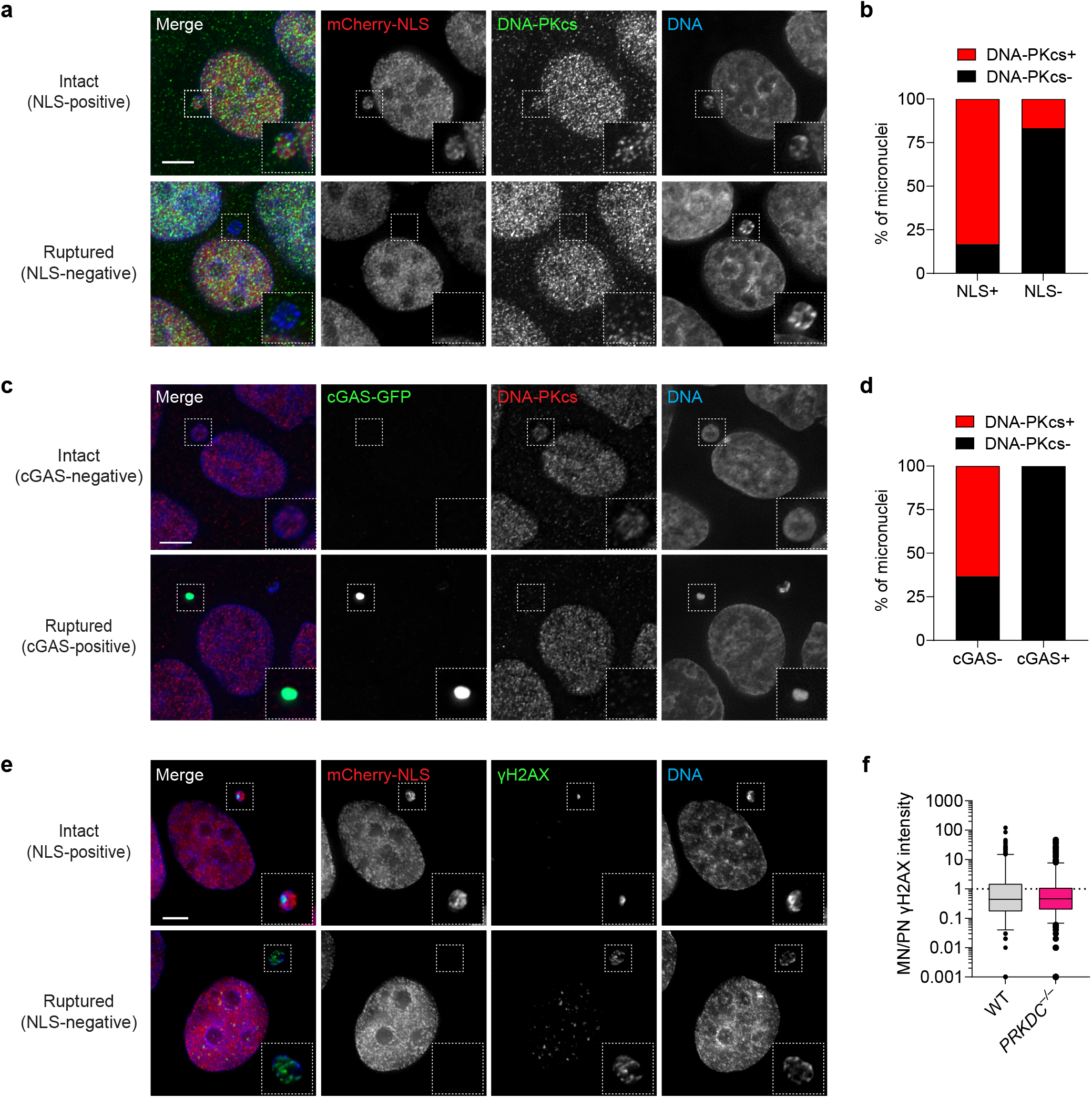
Nuclear envelope rupture triggers the loss of DNA-PKcs from micronuclei. **a)** Representative immunofluorescent images of DNA-PKcs in DLD-1 cells expressing mCherry fused to a nuclear localization signal (NLS) with intact (NLS-positive) or ruptured (NLS-negative) micronuclei. **b)** Proportion of intact and ruptured micronuclei with detectable levels of DNA-PKcs. Data represent *n* = 149 interphase cells with micronuclei. **c)** Representative immunofluorescent images of DNA-PKcs in DLD-1 cells expressing cGAS-GFP with intact (cGAS-negative) or ruptured (cGAS-positive) micronuclei. **d)** Proportion of intact and ruptured micronuclei with detectable levels of DNA-PKcs. Data represent *n* = 151 interphase cells with micronuclei. **e)** Representative immunofluorescent images of γH2AX in *PRKDC* KO DLD-1 cells expressing mCherry-NLS with intact or ruptured micronuclei. **f)** Fluorescence intensity of γH2AX in WT and *PRKDC* KO micronuclei compared to their corresponding primary nucleus. Data represent 10-90 percentile from *n* = 207 interphase cells with micronuclei for both WT and *PRKDC* KO. Scale bar for panels (**a**), (**c**), and (**e**), 5 µm.

**Extended Data Figure 7.**
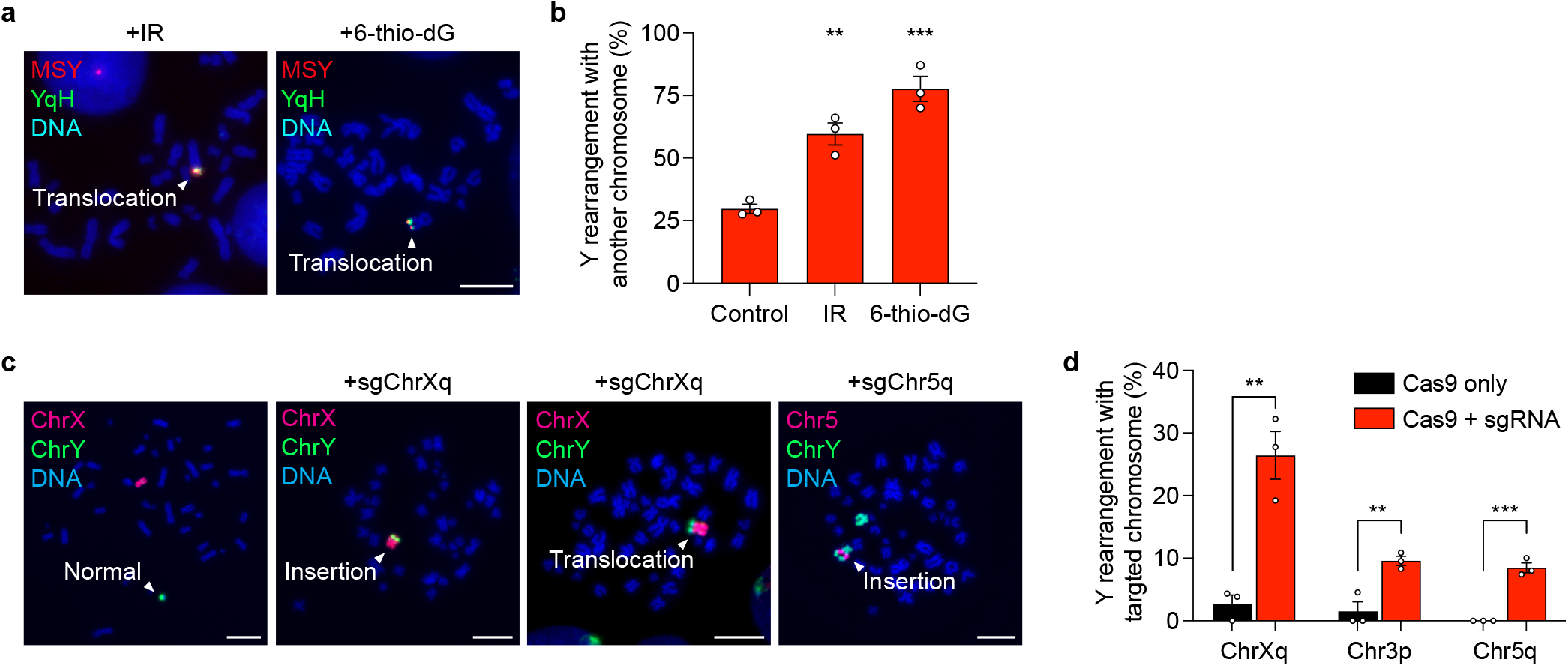
Induction of genome-wide or Cas9-induced DNA breaks on nuclear chromosomes facilitate translocations involving the micronucleated chromosome. **a)** Representative images of metaphase spreads hybridized to the indicated FISH probes showing examples of Y chromosome translocations. Scale bar, 10 µm. **b)** Frequency of inter-chromosomal rearrangements in cells following 3d DOX/IAA treatment, exposure to IR (2 Gy) or 6-thio-dG, and G418 selection. Data represent mean ± SEM of *n* = 3 independent experiments from (left to right): 222, 231, and 152 metaphase spreads with Y chromosome rearrangements. **c)** Representative images of metaphase spreads hybridized to the indicated FISH probes showing examples of normal or rearranged Y chromosomes with the indicated translocation partner. Scale bar, 10 µm. **d)** Frequency of Y chromosome rearrangements with the CRISPR/Cas9-targeted chromosome. Data represent mean ± SEM of *n* = 3 independent experiments from (left to right): 60, 75, 78,115, 84, and 77 metaphase spreads with Y chromosome rearrangements. Statistical analyses were calculated by ordinary one-way ANOVA test with multiple comparisons for panel (**b**) and by unpaired Student’s t-test for panel (**d**). \*\**P* ≤ 0.001; \*\*\**P* ≤ 0.001.

## SUPPLEMENTARY MATERIALS

**Supplementary Table 1.**
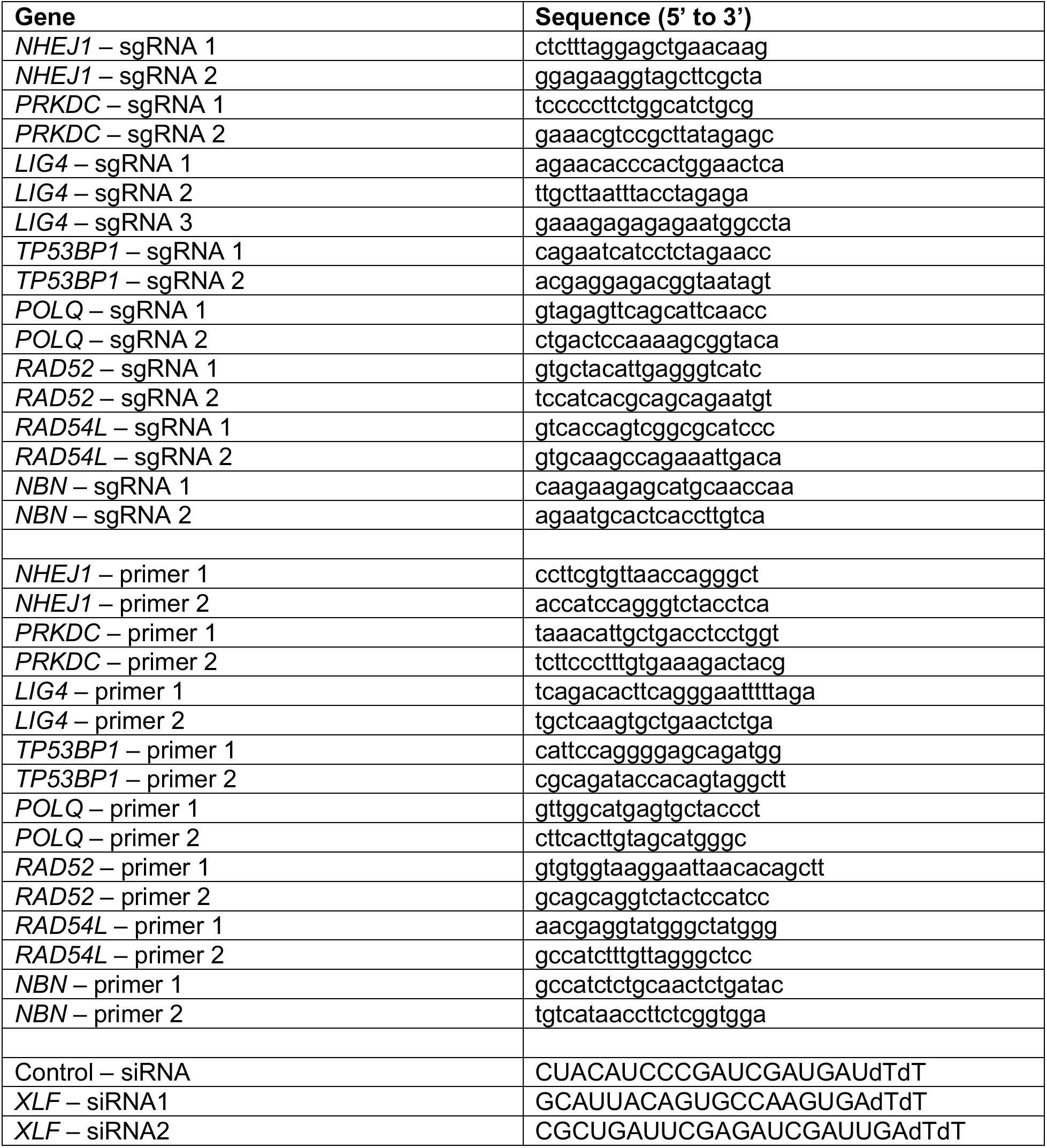
List of oligonucleotides sequences used in this study.

**Supplementary Table 2.** List of primary antibodies used in this study.

| **Antibody** | **Source** |
| --- | --- |
| XLF | Cell Signaling 2854 |
| DNA-PKcs | Abcam ab44815 (a gift from Benjamin Chen) |
| DNA Ligase IV | Cell Signaling 14649 |
| 53BP1 | Novus NB100-304 |
| RAD52 | Santa Cruz sc-365341 |
| RAD54 | Cell Signaling 15016 |
| NBS1 | Cell Signaling 14956 |
| α-Tubulin | Cell Signaling 3873 |
| Histone H2A.X (phospho S139) | Cell Signaling 2577 |
| Histone H2A.X (phospho S139) | Millipore 05-636 |
| FLAG | Sigma F1804 |
| GFP | Cell Signaling 2555 |
| mCherry | Cell Signaling 43590S |

## ACKNOWLEDGEMENTS

This work is dedicated to the memory of Benjamin Chen. We thank Anthony Davis for providing reagents and critical feedback; Benjamin Chen and Jerry Shay for providing reagents; Yu-Fen Lin and Haiyang Yu for technical assistance; and members of the Ly Laboratory for discussions. We acknowledge the UT Southwestern Flow Cytometry Core, Peter O’Donnell Jr. Brain Institute, and Department of Radiation Oncology for shared use of equipment. This work was supported by the US National Institutes of Health (R35GM146610 to P.L.), the Cancer Prevention and Research Institute of Texas (RR180050 to P.L.), and The Welch Foundation (I- 2071-20210327 to P.L.). J.E.V.-I. and I.C.-C. acknowledge the European Molecular Biology Laboratory for funding.

## Notes

https://www.ebi.ac.uk/ena/browser/home

### Competing Interest Statement

The authors have declared no competing interest.

## REFERENCES

1. Hatch EM, Fischer AH, Deerinck TJ, Hetzer MW. 2013. Catastrophic nuclear envelope collapse in cancer cell micronuclei. Cell 154:47–60.

2. Liu S, Kwon M, Mannino M, Yang N, Renda F, Khodjakov A, Pellman D. 2018. Nuclear envelope assembly defects link mitotic errors to chromothripsis. Nature 561:551–555.

3. Terradas M, Martín M, Tusell L, Genescà A. 2009. DNA lesions sequestered in micronuclei induce a local defective-damage response. DNA Repair 8:1225–1234.

4. Crasta K, Ganem NJ, Dagher R, Lantermann AB, Ivanova EV, Pan Y, Nezi L, Protopopov A, Chowdhury D, Pellman D. 2012. DNA breaks and chromosome pulverization from errors in mitosis. Nature 482:53–8.

5. Zhang CZ, Spektor A, Cornils H, Francis JM, Jackson EK, Liu S, Meyerson M, Pellman D. 2015. Chromothripsis from DNA damage in micronuclei. Nature 522:179–84.

6. Tang S, Stokasimov E, Cui Y, Pellman D. 2022. Breakage of cytoplasmic chromosomes by pathological DNA base excision repair. Nature 606:930–936.

7. Ly P, Teitz LS, Kim DH, Shoshani O, Skaletsky H, Fachinetti D, Page DC, Cleveland DW. 2017. Selective Y centromere inactivation triggers chromosome shattering in micronuclei and repair by non-homologous end joining. Nat Cell Biol 19:68–75.

8. Stephens PJ, Greenman CD, Fu B, Yang F, Bignell GR, Mudie LJ, Pleasance ED, Lau KW, Beare D, Stebbings LA, McLaren S, Lin ML, McBride DJ, Varela I, Nik-Zainal S, Leroy C, Jia M, Menzies A, Butler AP, Teague JW, Quail MA, Burton J, Swerdlow H, Carter NP, Morsberger LA, Iacobuzio-Donahue C, Follows GA, Green AR, Flanagan AM, Stratton MR, Futreal PA, Campbell PJ. 2011. Massive genomic rearrangement acquired in a single catastrophic event during cancer development. Cell 144:27–40.

9. Lin Y-F, Hu Q, Mazzagatti A, Valle-Inclán JE, Maurais EG, Dahiya R, Guyer A, Sanders JT, Engel JL, Nguyen G, Bronder D, Bakhoum SF, Cortés-Ciriano I, Ly P. 2023. Mitotic clustering of pulverized chromosomes from micronuclei. Nature 618:1041–1048.

10. Trivedi P, Steele CD, Au FKC, Alexandrov LB, Cleveland DW. 2023. Mitotic tethering enables inheritance of shattered micronuclear chromosomes. Nature 618:1049–1056.

11. Papathanasiou S, Mynhier NA, Liu S, Brunette G, Stokasimov E, Jacob E, Li L, Comenho C, van Steensel B, Buenrostro JD, Zhang CZ, Pellman D. 2023. Heritable transcriptional defects from aberrations of nuclear architecture. Nature 619:184–192.

12. Ly P, Cleveland DW. 2017. Rebuilding Chromosomes After Catastrophe: Emerging Mechanisms of Chromothripsis. Trends Cell Biol 27:917–930.

13. Cortés-Ciriano I, Lee JJ-K, Xi R, Jain D, Jung YL, Yang L, Gordenin D, Klimczak LJ, Zhang C-Z, Pellman DS, Akdemir KC, Alvarez EG, Baez-Ortega A, Beroukhim R, Boutros PC, Bowtell DDL, Brors B, Burns KH, Campbell PJ, Chan K, Chen K, Cortés-Ciriano I, Dueso-Barroso A, Dunford AJ, Edwards PA, Estivill X, Etemadmoghadam D, Feuerbach L, Fink JL, Frenkel-Morgenstern M, Garsed DW, Gerstein M, Gordenin DA, Haan D, Haber JE, Hess JM, Hutter B, Imielinski M, Jones DTW, Ju YS, Kazanov MD, Klimczak LJ, Koh Y, Korbel JO, Kumar K, Lee EA, Lee JJ-K, Li Y, Lynch AG, Macintyre G, et al. 2020. Comprehensive analysis of chromothripsis in 2,658 human cancers using whole-genome sequencing. Nature Genetics 52:331–341.

14. Voronina N, Wong JKL, Hubschmann D, Hlevnjak M, Uhrig S, Heilig CE, Horak P, Kreutzfeldt S, Mock A, Stenzinger A, Hutter B, Frohlich M, Brors B, Jahn A, Klink B, Gieldon L, Sieverling L, Feuerbach L, Chudasama P, Beck K, Kroiss M, Heining C, Mohrmann L, Fischer A, Schrock E, Glimm H, Zapatka M, Lichter P, Frohling S, Ernst A. 2020. The landscape of chromothripsis across adult cancer types. Nat Commun 11:2320.

15. Korbel JO, Campbell PJ. 2013. Criteria for inference of chromothripsis in cancer genomes. Cell 152:1226–36.

16. Ly P, Brunner SF, Shoshani O, Kim DH, Lan W, Pyntikova T, Flanagan AM, Behjati S, Page DC, Campbell PJ, Cleveland DW. 2019. Chromosome segregation errors generate a diverse spectrum of simple and complex genomic rearrangements. Nat Genet 51:705–715.

17. Scully R, Panday A, Elango R, Willis NA. 2019. DNA double-strand break repair-pathway choice in somatic mammalian cells. Nat Rev Mol Cell Biol doi:10.1038/s41580-019-0152-0.

18. Ramsden DA, Carvajal-Garcia J, Gupta GP. 2022. Mechanism, cellular functions and cancer roles of polymerase-theta-mediated DNA end joining. Nature Reviews Molecular Cell Biology 23:125–140.

19. Filippo JS, Sung P, Klein H. 2008. Mechanism of Eukaryotic Homologous Recombination. Annual Review of Biochemistry 77:229–257.

20. Bhargava R, Onyango DO, Stark JM. 2016. Regulation of Single-Strand Annealing and its Role in Genome Maintenance. Trends in Genetics 32:566–575.

21. Ferguson DO, Alt FW. 2001. DNA double strand break repair and chromosomal translocation: Lessons from animal models. Oncogene 20:5572–5579.

22. Stankiewicz P, Lupski JR. 2002. Genome architecture, rearrangements and genomic disorders. Trends in Genetics 18:74–82.

23. Kasparek TR, Humphrey TC. 2011. DNA double-strand break repair pathways, chromosomal rearrangements and cancer. Seminars in Cell & Developmental Biology 22:886–897.

24. Ceccaldi R, Rondinelli B, D’Andrea AD. 2016. Repair Pathway Choices and Consequences at the Double-Strand Break. Trends in Cell Biology 26:52–64.

25. Piazza A, Wright WD, Heyer WD. 2017. Multi-invasions Are Recombination Byproducts that Induce Chromosomal Rearrangements. Cell 170:760–773 e15.

26. Kloosterman WP, Guryev V, van Roosmalen M, Duran KJ, de Bruijn E, Bakker SC, Letteboer T, van Nesselrooij B, Hochstenbach R, Poot M, Cuppen E. 2011. Chromothripsis as a mechanism driving complex de novo structural rearrangements in the germline. Hum Mol Genet 20:1916–24.

27. Kloosterman WP, Hoogstraat M, Paling O, Tavakoli-Yaraki M, Renkens I, Vermaat JS, van Roosmalen MJ, van Lieshout S, Nijman IJ, Roessingh W, van ’t Slot R, van de Belt J, Guryev V, Koudijs M, Voest E, Cuppen E. 2011. Chromothripsis is a common mechanism driving genomic rearrangements in primary and metastatic colorectal cancer. Genome Biol 12:R103.

28. Chiang C, Jacobsen JC, Ernst C, Hanscom C, Heilbut A, Blumenthal I, Mills RE, Kirby A, Lindgren AM, Rudiger SR, McLaughlan CJ, Bawden CS, Reid SJ, Faull RL, Snell RG, Hall IM, Shen Y, Ohsumi TK, Borowsky ML, Daly MJ, Lee C, Morton CC, MacDonald ME, Gusella JF, Talkowski ME. 2012. Complex reorganization and predominant non-homologous repair following chromosomal breakage in karyotypically balanced germline rearrangements and transgenic integration. Nat Genet 44:390–7, S1.

29. Boeva V, Jouannet S, Daveau R, Combaret V, Pierre-Eugene C, Cazes A, Louis-Brennetot C, Schleiermacher G, Ferrand S, Pierron G, Lermine A, Rio Frio T, Raynal V, Vassal G, Barillot E, Delattre O, Janoueix-Lerosey I. 2013. Breakpoint features of genomic rearrangements in neuroblastoma with unbalanced translocations and chromothripsis. PLoS One 8:e72182.

30. Weckselblatt B, Hermetz KE, Rudd MK. 2015. Unbalanced translocations arise from diverse mutational mechanisms including chromothripsis. Genome Res 25:937–47.

31. Tan EH, Henry IM, Ravi M, Bradnam KR, Mandakova T, Marimuthu MP, Korf I, Lysak MA, Comai L, Chan SW. 2015. Catastrophic chromosomal restructuring during genome elimination in plants. Elife 4.

32. Ratnaparkhe M, Wong JKL, Wei P-C, Hlevnjak M, Kolb T, Simovic M, Haag D, Paul Y, Devens F, Northcott P, Jones DTW, Kool M, Jauch A, Pastorczak A, Mlynarski W, Korshunov A, Kumar R, Downing SM, Pfister SM, Zapatka M, McKinnon PJ, Alt FW, Lichter P, Ernst A. 2018. Defective DNA damage repair leads to frequent catastrophic genomic events in murine and human tumors. Nature Communications 9:4760.

33. Cleal K, Jones RE, Grimstead JW, Hendrickson EA, Baird DM. 2019. Chromothripsis during telomere crisis is independent of NHEJ, and consistent with a replicative origin. Genome Research 29:737–749.

34. Symington L, Gautier J. 2011. Double-strand break end resection and repair pathway choice. Annual review of genetics 45:247–71.

35. Maciejowski J, Li Y, Bosco N, Campbell Peter J, de Lange T. 2015. Chromothripsis and Kataegis Induced by Telomere Crisis. Cell 163:1641–1654.

36. Kurimasa A, Kumano S, Boubnov NV, Story MD, Tung C-S, Peterson SR, Chen DJ. 1999. Requirement for the Kinase Activity of Human DNA-Dependent Protein Kinase Catalytic Subunit in DNA Strand Break Rejoining. Molecular and Cellular Biology 19:3877–3884.

37. Chaplin AK, Hardwick SW, Stavridi AK, Buehl CJ, Goff NJ, Ropars V, Liang S, De Oliveira TM, Chirgadze DY, Meek K, Charbonnier J-B, Blundell TL. 2021. Cryo-EM of NHEJ supercomplexes provides insights into DNA repair. Molecular Cell 81:3400–3409.e3.

38. Chen X, Xu X, Chen Y, Cheung JC, Wang H, Jiang J, de Val N, Fox T, Gellert M, Yang W. 2021. Structure of an activated DNA-PK and its implications for NHEJ. Molecular Cell 81:801–810.e3.

39. Scott DE, Francis-Newton NJ, Marsh ME, Coyne AG, Fischer G, Moschetti T, Bayly AR, Sharpe TD, Haas KT, Barber L, Valenzano CR, Srinivasan R, Huggins DJ, Lee M, Emery A, Hardwick B, Ehebauer M, Dagostin C, Esposito A, Pellegrini L, Perrior T, McKenzie G, Blundell TL, Hyvönen M, Skidmore J, Venkitaraman AR, Abell C. 2021. A small-molecule inhibitor of the BRCA2-RAD51 interaction modulates RAD51 assembly and potentiates DNA damage-induced cell death. Cell Chemical Biology 28:835–847.e5.

40. Zatreanu D, Robinson HMR, Alkhatib O, Boursier M, Finch H, Geo L, Grande D, Grinkevich V, Heald RA, Langdon S, Majithiya J, McWhirter C, Martin NMB, Moore S, Neves J, Rajendra E, Ranzani M, Schaedler T, Stockley M, Wiggins K, Brough R, Sridhar S, Gulati A, Shao N, Badder LM, Novo D, Knight EG, Marlow R, Haider S, Callen E, Hewitt G, Schimmel J, Prevo R, Alli C, Ferdinand A, Bell C, Blencowe P, Bot C, Calder M, Charles M, Curry J, Ekwuru T, Ewings K, Krajewski W, MacDonald E, McCarron H, Pang L, Pedder C, Rigoreau L, Swarbrick M, et al. 2021. Polθ inhibitors elicit BRCA-gene synthetic lethality and target PARP inhibitor resistance. Nature Communications 12:3636.

41. Menear KA, Adcock C, Boulter R, Cockcroft XL, Copsey L, Cranston A, Dillon KJ, Drzewiecki J, Garman S, Gomez S, Javaid H, Kerrigan F, Knights C, Lau A, Loh VM, Jr., Matthews IT, Moore S, O’Connor MJ, Smith GC, Martin NM. 2008. 4-[3-(4- cyclopropanecarbonylpiperazine-1-carbonyl)-4-fluorobenzyl]-2H-phthalazin-1-one: a novel bioavailable inhibitor of poly(ADP-ribose) polymerase-1. J Med Chem 51:6581–91.

42. Chen S, Lee L, Naila T, Fishbain S, Wang A, Tomkinson AE, Lees-Miller SP, He Y. 2021. Structural basis of long-range to short-range synaptic transition in NHEJ. Nature 593:294–298.

43. Oksenych V, Kumar V, Liu X, Guo C, Schwer B, Zha S, Alt FW. 2013. Functional redundancy between the XLF and DNA-PKcs DNA repair factors in V(D)J recombination and nonhomologous DNA end joining. Proc Natl Acad Sci U S A 110:2234–9.

44. Cisneros-Aguirre M, Lopezcolorado FW, Tsai LJ, Bhargava R, Stark JM. 2022. The importance of DNAPKcs for blunt DNA end joining is magnified when XLF is weakened. Nature Communications 13:3662.

45. Mackenzie KJ, Carroll P, Martin C-A, Murina O, Fluteau A, Simpson DJ, Olova N, Sutcliffe H, Rainger JK, Leitch A, Osborn RT, Wheeler AP, Nowotny M, Gilbert N, Chandra T, Reijns MAM, Jackson AP. 2017. cGAS surveillance of micronuclei links genome instability to innate immunity. Nature 548:461–465.

46. Harding SM, Benci JL, Irianto J, Discher DE, Minn AJ, Greenberg RA. 2017. Mitotic progression following DNA damage enables pattern recognition within micronuclei. Nature 548:466–470.

47. Terasawa M, Shinohara A, Shinohara M. 2014. Canonical non-homologous end joining in mitosis induces genome instability and is suppressed by M-phase-specific phosphorylation of XRCC4. PLoS Genet 10:e1004563.

48. Zgheib O, Pataky K, Brugger J, Halazonetis TD. 2009. An oligomerized 53BP1 tudor domain suffices for recognition of DNA double-strand breaks. Mol Cell Biol 29:1050–8.

49. Fok JHL, Ramos-Montoya A, Vazquez-Chantada M, Wijnhoven PWG, Follia V, James N, Farrington PM, Karmokar A, Willis SE, Cairns J, Nikkilä J, Beattie D, Lamont GM, Finlay MRV, Wilson J, Smith A, O’Connor LO, Ling S, Fawell SE, O’Connor MJ, Hollingsworth SJ, Dean E, Goldberg FW, Davies BR, Cadogan EB. 2019. AZD7648 is a potent and selective DNA-PK inhibitor that enhances radiation, chemotherapy and olaparib activity. Nature Communications 10:5065.

50. Mender I, Gryaznov S, Dikmen ZG, Wright WE, Shay JW. 2015. Induction of Telomere Dysfunction Mediated by the Telomerase Substrate Precursor 6-Thio-2′- Deoxyguanosine. Cancer Discovery 5:82–95.

51. Mohr L, Toufektchan E, von Morgen P, Chu K, Kapoor A, Maciejowski J. 2021. ER-directed TREX1 limits cGAS activation at micronuclei. Molecular Cell 81:724–738.e9.

52. Umbreit NT, Zhang C-Z, Lynch LD, Blaine LJ, Cheng AM, Tourdot R, Sun L, Almubarak HF, Judge K, Mitchell TJ, Spektor A, Pellman D. 2020. Mechanisms generating cancer genome complexity from a single cell division error. Science 368:eaba0712.

53. Hastings PJ, Ira G, Lupski JR. 2009. A microhomology-mediated break-induced replication model for the origin of human copy number variation. PLoS Genet 5:e1000327.

54. Sakofsky CJ, Ayyar S, Deem AK, Chung WH, Ira G, Malkova A. 2015. Translesion Polymerases Drive Microhomology-Mediated Break-Induced Replication Leading to Complex Chromosomal Rearrangements. Mol Cell 60:860–72.

55. Deng L, Wu RA, Sonneville R, Kochenova OV, Labib K, Pellman D, Walter JC. 2019. Mitotic CDK Promotes Replisome Disassembly, Fork Breakage, and Complex DNA Rearrangements. Molecular Cell 73:915–929.e6.

56. Gelot C, Kovacs MT, Miron S, Mylne E, Ghouil R, Popova T, Dingli F, Loew D, Guirouilh-Barbat J, Nery ED, Zinn-Justin S, Ceccaldi R. 2023. Polθ is phosphorylated by Polo-like kinase 1 (PLK1) to enable repair of DNA double strand breaks in mitosis. bioRxiv doi:10.1101/2023.03.17.533134:2023.03.17.533134.

57. Brambati A, Sacco O, Porcella S, Heyza J, Kareh M, Schmidt JC, Sfeir A. 2023. RHINO directs MMEJ to repair DNA breaks in mitosis. Science doi:10.1126/science.adh3694:eadh3694.

58. Janssen A, van der Burg M, Szuhai K, Kops GJPL, Medema RH. 2011. Chromosome Segregation Errors as a Cause of DNA Damage and Structural Chromosome Aberrations. Science 333:1895–1898.

59. Shoshani O, Brunner SF, Yaeger R, Ly P, Nechemia-Arbely Y, Kim DH, Fang R, Castillon GA, Yu M, Li JSZ, Sun Y, Ellisman MH, Ren B, Campbell PJ, Cleveland DW. 2021. Chromothripsis drives the evolution of gene amplification in cancer. Nature 591:137–141.

60. Dharanipragada P, Zhang X, Liu S, Lomeli SH, Hong A, Wang Y, Yang Z, Lo KZ, Vega-Crespo A, Ribas A, Moschos SJ, Moriceau G, Lo RS. 2023. Blocking Genomic Instability Prevents Acquired Resistance to MAPK Inhibitor Therapy in Melanoma. Cancer Discov 13:880–909.

61. Li H, Durbin R. 2010. Fast and accurate long-read alignment with Burrows-Wheeler transform. Bioinformatics 26:589–95.

62. Danecek P, Bonfield JK, Liddle J, Marshall J, Ohan V, Pollard MO, Whitwham A, Keane T, McCarthy SA, Davies RM, Li H. 2021. Twelve years of SAMtools and BCFtools. Gigascience 10.

63. Tarasov A, Vilella AJ, Cuppen E, Nijman IJ, Prins P. 2015. Sambamba: fast processing of NGS alignment formats. Bioinformatics 31:2032–4.

64. Pedersen BS, Quinlan AR. 2018. Mosdepth: quick coverage calculation for genomes and exomes. Bioinformatics 34:867–868.

65. Boeva V, Popova T, Bleakley K, Chiche P, Cappo J, Schleiermacher G, Janoueix-Lerosey I, Delattre O, Barillot E. 2012. Control-FREEC: a tool for assessing copy number and allelic content using next-generation sequencing data. Bioinformatics 28:423–5.

66. Manders F, Brandsma AM, de Kanter J, Verheul M, Oka R, van Roosmalen MJ, van der Roest B, van Hoeck A, Cuppen E, van Boxtel R. 2022. MutationalPatterns: the one stop shop for the analysis of mutational processes. BMC Genomics 23:134.

